# Modulation of Tau Tubulin Kinases (TTBK1 and TTBK2) Impacts Ciliogenesis

**DOI:** 10.1101/2022.05.06.490937

**Authors:** Frances M. Potjewyd, Ariana B. Marquez, Apirat Chaikuad, Stefanie Howell, Andrea S. Dunn, Alvaro A. Beltran, Jeffery L. Smith, David H. Drewry, Adriana S. Beltran, Alison D. Axtman

## Abstract

Tau tubulin kinase 1 and 2 (TTBK1 and TTBK2) are highly homologous kinases that are expressed and mediate disease-relevant pathways predominantly in the brain. Distinct roles for TTBK1 and TTBK2 have been delineated. While efforts have been devoted to characterizing the impact to TTBK1 inhibition in diseases like Alzheimer’s disease and amyotrophic lateral sclerosis, TTBK2 inhibition has been less explored. TTBK2 serves a critical function during cilia assembly. Given the biological importance of these kinases, we designed a targeted library from which we identified several chemical tools that engage TTBK1 and TTBK2 in cells and inhibit their downstream signaling. Indolyl pyrimidinamine 10 significantly reduced the expression of primary cilia on the surface of human induced pluripotent stem cells (iPSCs). Furthermore, analog 10 phenocopies TTBK2 KO in iPSCs, confirming an essential role for TTBK2 in ciliogenesis.

## INTRODUCTION

Tau tubulin kinase 1 and 2 (TTBK1 and TTBK2) are two understudied serine/threonine/tyrosine kinases that belong to the casein kinase 1 superfamily. The kinase domains of TTBK1 and TTBK2 have 88% identity and 96% similarity and the same catalytic residues: lysine 63 and aspartic acid 164 for TTBK1 and lysine 50 and aspartic acid 141 for TTBK2 (Nozal and Martinez, 2019). The non-catalytic domains of these enzymes are distinct from one another (Ikezu and Ikezu, 2014). Several apo and co-crystal structures have been solved and deposited in the PDB for these proteins, including seven structures for human TTBK1 and three structures for human TTBK2. In addition to phosphorylation of several characterized substrates, distinct physiological roles for these kinases have been described in key processes such as mitosis, ciliogenesis, microtubule dynamics, glucose and GABA transport, neurotransmission, and neuroinflammation (Ikezu and Ikezu, 2014; Liao et al., 2015; Nozal and Martinez, 2019).

Expression of TTBK1 is localized to the brain, especially in the cytoplasm of cortical, hippocampal, and entorhinal cortex neurons, and overexpressed in neurodegenerative diseases (Ikezu et al., 2020; Nozal and Martinez, 2019; Sato et al., 2006). Although TTBK2 is ubiquitously expressed in human tissues, higher expression has been reported in the cerebellum Purkinje cells, granular cell layer, hippocampus, midbrain, and substantia nigra (Houlden et al., 2007). Consistent with these expression patterns, these kinases also have described pathological roles, including many specific to the central nervous system and brain such as Alzheimer’s disease (AD), frontotemporal dementia (FTD), amyotrophic lateral sclerosis (ALS), and a spinocerebellar ataxia (McMillan et al., 2020). A list of distinct neuronal interactors and phosphorylation substrates for TTBK1 and TTBK2 has been generated to address the implications of potential therapeutic targeting of each (Bao et al., 2021).

As their name implies, these kinases are reported to phosphorylate tubulin and tau at multiple serine residues. These same epitopes are also hyperphosphorylated in neurofibrillary tangles (NFT), a key pathological hallmark in several neurodegenerative diseases, including AD (Sato *et al*., 2006). Furthermore, TTBK1 expression induces neurite and axonal degeneration, indicating other critical functions for this kinase in early AD pathology (Ikezu and Ikezu, 2014; Ikezu *et al*., 2020). In addition to these roles in early AD, TTBK1 is upregulated in the frontal cortex of AD patients and single nucleotide polymorphisms (SNPs) in the TTBK1 gene are associated with late-onset AD (LOAD) (Sato et al., 2008; Yu et al., 2011). Specific polymorphisms were found to decrease the risk of LOAD in multiple patient populations (Yu *et al*., 2011). Abundant expression of TTBK1/2 in the brain coupled with their confirmed role of phosphorylating tau and leading to NFT supports the hypothesis that TTBK1/2 inhibitors may be therapeutically beneficial to AD patients (Dillon et al., 2020; Lund et al., 2013).

With respect to ALS and FTD, TTBK1 and TTBK2 phosphorylate TAR DNA binding protein of 43 kDa (TDP-43) both *in vitro* and *in vivo* (Liachko et al., 2014). This phosphorylation causes TDP-43 precipitation and aggregation in the cytoplasm of neurons, contributing to toxicity and eventual neuronal death (Nozal and Martinez, 2019). Furthermore, these kinases co-localize with TDP-43 inclusions in the frontal cortex and spinal cord of FTD and ALS patient, respectively (Liachko *et al*., 2014; McMillan *et al*., 2020; Nozal and Martinez, 2019). Reduction of TTBK1 mRNA decreased the levels of phosphorylated TDP-43 in a pathological model, suggesting that inhibitors could also provide relief for FTD/ALS patients.

A pathogenic variant in the TTBK2 gene that results in premature truncations of the corresponding protein causes a rare neurological disorder called spinocerebellar ataxia type 11 (SCA11) (Bowie et al., 2018; Houlden *et al*., 2007; Nozal and Martinez, 2019). SCA11 is characterized by atrophy of Purkinje neurons in the cerebellum (Bowie *et al*., 2018). Interestingly, inducible TTBK1 transgenic mice provide a model of cerebellar neurodegeneration with phenotypes reminiscent of spinocerebellar ataxia (McMillan *et al*., 2020). SCA11 alleles of TTBK2 are reported to interfere with ciliogenesis and cilium stability (Bowie *et al*., 2018). Accordingly, several centriolar and distal end proteins are substrates of TTBK2, recruitment of TTBK2 enables requisite steps of cilia assembly, and this kinase is essential for regulating the growth of axonemal microtubules during ciliogenesis (Liao *et al*., 2015; May et al., 2021). Loss of TTBK2 has been reported to permit basal body docking to the plasma membrane while impeding transition zone formation and ciliary shaft elongation (May *et al*., 2021). Moreover, emerging studies suggest that abnormalities in the length and frequency of primary cilia could be a key regulator in brain diseases (Ma et al., 2022).

Our program directed at targeting TTBK1/2 was initiated based upon two complementary motivations. First, we are interested in the generation and use of kinase inhibitors as tools to increase understanding of signaling pathways that drive neurodegenerative disease. As outlined in this section, TTBK1 and TTBK2 have several described functions in the brain and have accordingly become sought after targets for neurodegenerative disease therapies. Second, we seek to identify high-quality tool molecules for understudied kinase targets. Both TTBK1 and TTBK2 were identified as “dark” kinases by the NIH Illuminating the Druggable Genome (IDG) program. The IDG program is accelerating characterization of the proteome via stimulation of research around those proteins that are most poorly studied, which they refer to as “dark” (Rodgers et al., 2018). We are supported by the arm of the IDG program targeting illumination of the dark kinome and are doing so via concurrent development of high-quality chemical tools and novel cell-based kinase assays for these understudied kinases.

Interest in the development of TTBK1/2 inhibitors has recently increased. Co-crystal structures have been deposited for two AstraZeneca inhibitors (AZ1 and AZ2, Figure 1A) and one inhibitor discovered by Bristol-Myers Squibb (BMS1, Figure 1A) (Xue et al., 2013). A recent paper described optimization of the AZ2 scaffold to yield compound 29 (Figure 1A), which demonstrated good brain penetration and was able to reduce TDP-43 phosphorylation *in vivo* (Nozal et al., 2022). Biogen also solved the only co-crystal structure of TTBK2 bound to an inhibitor (AZ1, Figure 1A) (Marcotte et al., 2020). A Biogen-led effort to deliver a potent and selective TTBK inhibitor with suitable CNS penetration for *in vivo* pharmacology studies resulted in preparation of BGN8, BGN18, and BGN31 (Figure 1A) as well as a co-crystal structure of BGN18 bound to TTBK1. These inhibitors were developed to test hypotheses related to modulation of tau phosphorylation in hypothermia and developmental animal models (Halkina et al., 2021). BGN31 (also called BIIB-TTBKi and TTBK1-IN-1) was shown to significantly lower tau phosphorylation at multiple disease-relevant sites when dosed *in vivo* (Dillon *et al*., 2020). It is worth noting that only for BGN18 and BGN31 has extensive selectivity profiling data been reported (Halkina *et al*., 2021).

**Figure 1.**
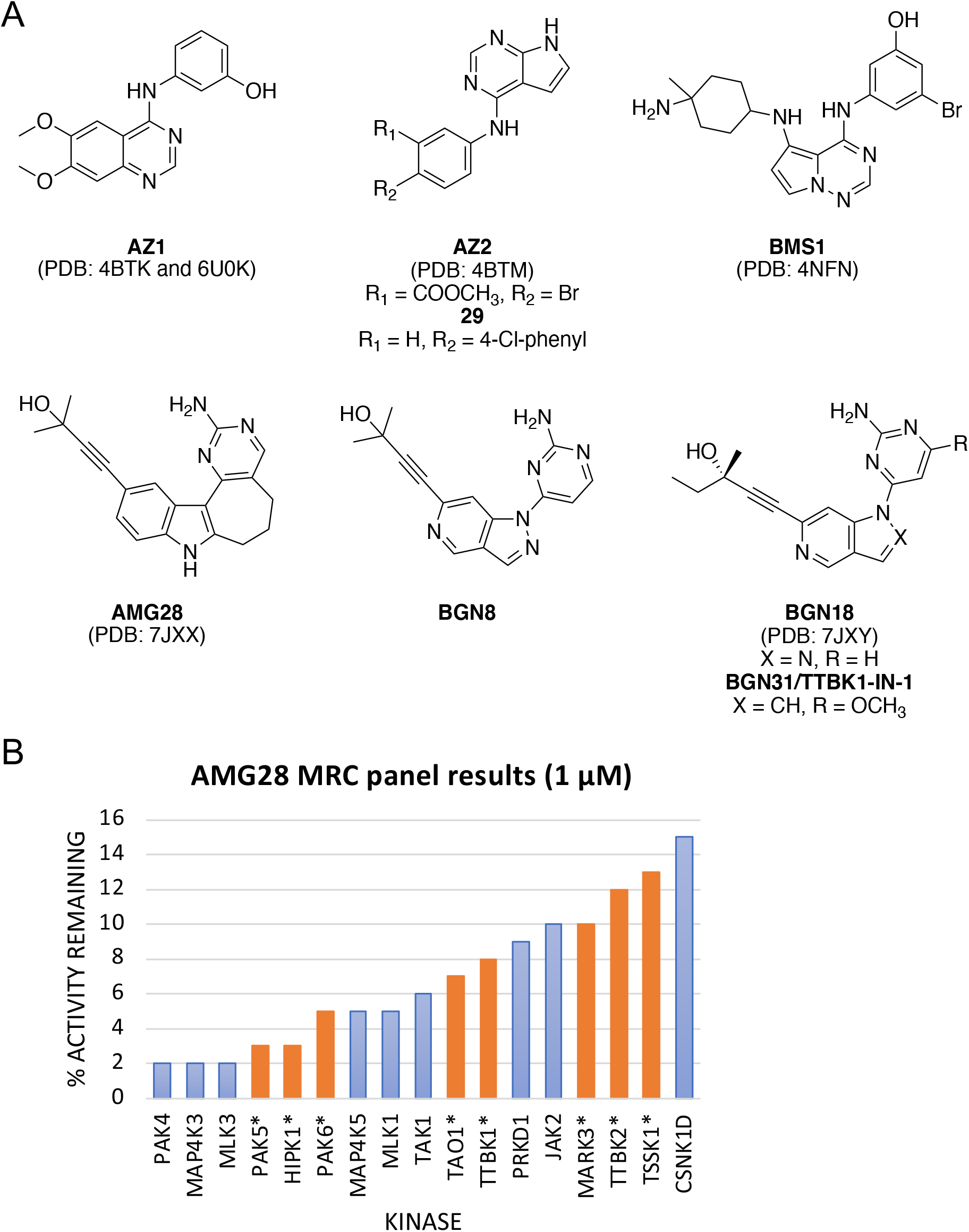
Structures and data for published TTBK inhibitors. (A) Structures of published TTBK1/2 inhibitors and PDB codes for corresponding structures. (B) AMG28 MRC panel screening results for kinases with <15% activity remaining. IDG kinases marked with asterisk and orange bars.

Commercially available assays for TTBK1 and TTBK2 are limited. A 2017 survey of biochemical kinase assays offered by the largest vendors identified that only half had available TTBK1 and TTBK2 assays (Drewry et al., 2017). These kinases are not included in the largest kinome-wide profiling panel (Eurofins DiscoverX *scan*MAX), and thus the activity of broadly profiled compounds on TTBK1/2 could be overlooked. One panel that does include TTBK1 and TTBK2 is that offered by the International Centre for Kinase profiling within the MRC Protein Phosphorylation Unit at the University of Dundee (https://www.kinase-screen.mrc.ac.uk/kinase-inhibitors). However, the translation of activity in these biochemical assays to a cellular context is unknown, and there is a need for target-specific cell-based assays to support medicinal chemistry optimization efforts aimed at delivering cell-active inhibitors.

The high homology of the kinase domains of TTBK1 and TTBK2 makes it difficult to achieve selectivity between the two kinases. This is true for the compounds included in Figure 1A, which display nearly equal potency for both enzymes and are often described as TTBK inhibitors for this reason. This selectivity complication aside, the most advanced inhibitors, developed by Biogen, have been used to explore biology mediated by TTBK1 (Dillon *et al*., 2020; Halkina *et al*., 2021). Our efforts have been dedicated to exploring a process ascribed to TTBK2: ciliogenesis. We have developed high-quality, cell-active, small molecule dual inhibitors of TTBK1/2 and illustrated their impact on ciliogenesis.

## RESULTS

### Design and Synthesis of Indolyl Pyrimidinamine Inhibitors of TTBK1/2

Like the Biogen lead optimization effort, we took our lead (AMG28, Figure 1A) from examination of the off-target activity of a published Amgen NF-κB inducing kinase (NIK) inhibitor (Li et al., 2013). The potency of this compound was reported in the MRC Kinase Profiling Inhibitor Database as 8% activity remaining for TTBK1 and 12% activity remaining for TTBK2 when screened at 1 μM (Figure 1B) (https://www.kinase-screen.mrc.ac.uk/kinase-inhibitors). Biogen scientists evaluated this compound in a biochemical assay (TTBK1 IC_50_ = 199 nM) and in a cellular assay that measures inhibition of tau phosphorylation at Ser422 (IC_50_ = 1.85 μM). Like us (Figure 2A), this group solved a co-crystal structure of AMG28 with human TTBK1 (kinase domain only) and confirmed hinge-binding of the aminopyrimidine ring and that the alkyne displaces the catalytic lysine to project its tertiary alcohol into the back pocket. Furthermore, they determined that the N-H bond of the indole is solvent exposed and the seven-membered ring helps maintain the planarity of the molecule (Halkina *et al*., 2021).

**Figure 2.**
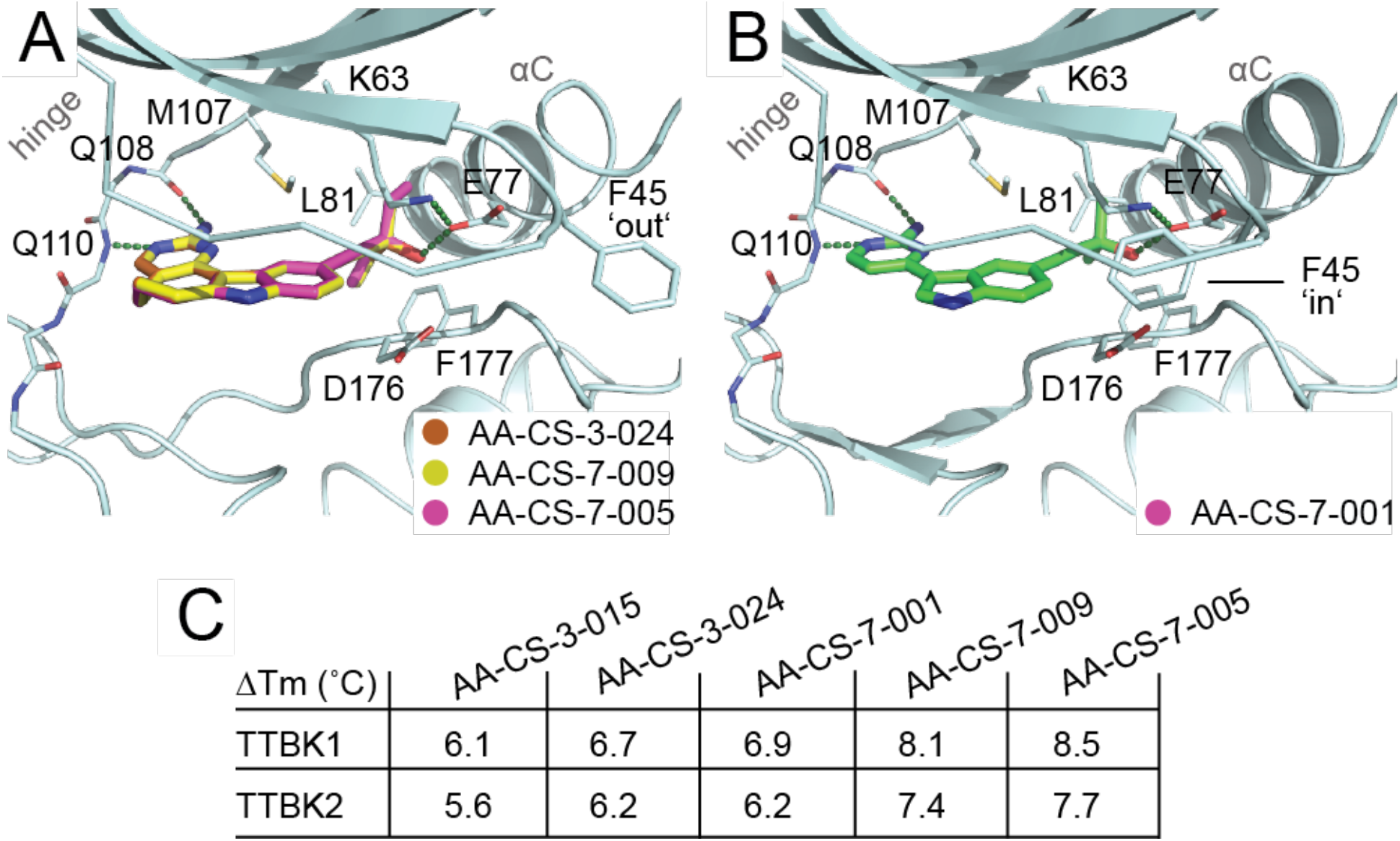
X-ray crystallographic structure of human TTBK1 in complex with several analogs and corresponding thermal shift data for TTBK1 and TTBK2. (A) AMG28 (PDB code: 7ZHN) is shown in brown, **10** (PDB code: 7ZHQ) in yellow, and **9** (PDB code: 7ZHP) in pink stick representation, respectively. (B) **3** (PDB code: 7ZHO) is shown in pink. The hinge region and αC helix are labeled in each panel and hydrogen bonds are included as black dashed lines. (C) Table of thermal shift (ΔTm) data for TTBK inhibitors.

Our first analogs were dedicated to understanding the role that the saturated ring system plays on TTBK1/2 activity. We hypothesized that the rotational freedom of AMG28 is reduced due to incorporation of a seven-membered ring as part of its framework and that it also helps project the alkynyl side chain into the back pocket. To investigate whether smaller or larger rings change the activity of this scaffold, we made the corresponding analogs that bear a six- and eight-membered ring, **1** and **2**, respectively. In addition, we removed the ring to make corresponding analog **3**. These analogs maintained the substitution of AMG28 and only varied in the ring size or absence of a ring.

Based on the potency of AMG28, we simultaneously explored changes to the substituent on the alkyne while retaining the seven-membered ring core scaffold. The free alcohol was unmodified in almost all analogs, with the exception of **11** in which the alcohol was capped as a methoxy group. Analog **11** was designed to determine whether that hydrogen on the alcohol is important for hydrogen bonding. Other analogs sought to probe whether changing the nature of the groups attached to the carbon center bearing the alcohol to reduce the steric complexity (**4** and **5**) or increase their length from methyl to ethyl (**9** and **10**) is beneficial. Finally, the remaining three analogs (**6**–**8**) projected the alcohol further into the pocket by adding another methylene unit to the chain. These analogs were designed to vary in steric bulk around the alcohol to simultaneously explore the tolerance of this deeper part of the pocket to alkyl substitution.

### Selective Enzymatic Kinase Inhibition Screening Indicates Inhibition of TTBK1 and TTBK2 by Several Indolyl Pyrimidinamine Analogs

Once prepared, we screened our library in a 17-member panel of enzymatic kinase assays (Table S1). Kinases were selected based on the MRC panel screening results for AMG28 with a bias toward the inclusion of understudied IDG kinases, a designation annotated in Table S1 and in Figure 1B for 14 of the 17 kinases (https://www.kinase-screen.mrc.ac.uk/kinase-inhibitors). A range of understudied kinases were potently inhibited by AMG28 within this MRC panel. Small families of kinases were chosen based on the availability of enzymatic assays offered by Eurofins. In addition to TTBK1 and TTBK2, PAK3–6, TAOK1–3, TSSK1–4, and MARK3–4 were included based on potent inhibition of one or more kinase(s) in the family by AMG28 when profiled at 1 μM (Figure 1B). Furthermore, PAK, MARK, and TAOK family kinases are of interest since they have implications in neurodegenerative disease signaling (Drewry et al., 2022a). The addition of HIPK1 and MAP4K5 completed our panel.

We profiled all 12 compounds (**1**–**11** and AMG28) against this kinase panel at a single concentration (1 μM) in radiometric enzymatic assays at the K_m_ = ATP for each kinase. Table S1 contains the results from this screen reported as percent of control (PoC) for each analog for each kinase, with lower values indicative of greater inhibition. A subset of this data is also collated in Table 1. PoC data for both TTBK1 and TTBK2 is included in Table 1 for each compound. The column labeled “# <10 PoC in enzyme panel” in Table 1 reports the number of kinases in this 17-member panel inhibited >90% by each analog.

**Table 1.**
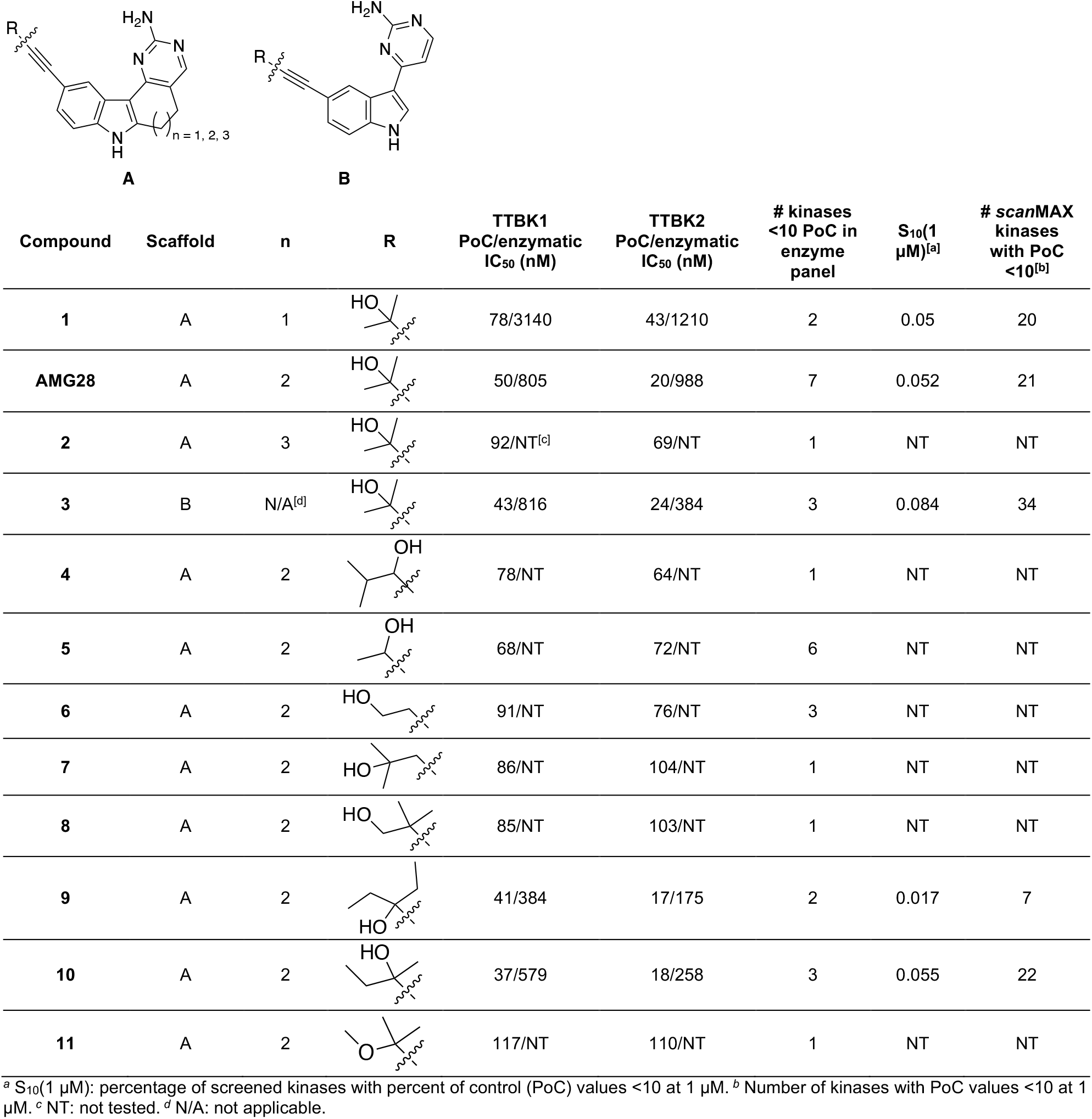
Kinase Panel Profiling of Indolyl Pyrimidinamine Library.

|  |  |  |  |  |  |  |  |  |
| --- | --- | --- | --- | --- | --- | --- | --- | --- |
| Compound | Scaffold | n | R | TTBK1<br>PoC/enzymatic<br>IC<sub>50</sub> (nM) | TTBK2<br>PoC/enzymatic<br>IC<sub>50</sub> (nM) | # kinases<br><10 PoC in<br>enzyme<br>panel | S<sub>10</sub>(1<br>μM)<sup>[a]</sup> | # scanMAX<br>kinases<br>with PoC<br><10<sup>[b]</sup> |
| 1 | A | 1 |  | 78/3140 | 43/1210 | 2 | 0.05 | 20 |
| AMG28 | A | 2 |  | 50/805 | 20/988 | 7 | 0.052 | 21 |
| 2 | A | 3 |  | 92/NT<sup>[c]</sup> | 69/NT | 1 | NT | NT |
| 3 | B | N/A<sup>[d]</sup> |  | 43/816 | 24/384 | 3 | 0.084 | 34 |
| 4 | A | 2 |  | 78/NT | 64/NT | 1 | NT | NT |
| 5 | A | 2 |  | 68/NT | 72/NT | 6 | NT | NT |
| 6 | A | 2 |  | 91/NT | 76/NT | 3 | NT | NT |
| 7 | A | 2 |  | 86/NT | 104/NT | 1 | NT | NT |
| 8 | A | 2 |  | 85/NT | 103/NT | 1 | NT | NT |
| 9 | A | 2 |  | 41/384 | 17/175 | 2 | 0.017 | 7 |
| 10 | A | 2 |  | 37/579 | 18/258 | 3 | 0.055 | 22 |
| 11 | A | 2 |  | 117/NT | 110/NT | 1 | NT | NT |

<sup>a</sup> S<sub>10</sub>(1 μM): percentage of screened kinases with percent of control (PoC) values <10 at 1 μM. <sup>b</sup> Number of kinases with PoC values <10 at 1 μM. <sup>c</sup> NT: not tested. <sup>d</sup> N/A: not applicable.

This single-concentration data provided an initial indication of potency for TTBK1 and/or TTBK2 as well as selectivity versus off-targets of parent molecule AMG28. Gratifyingly, we identified analogs of AMG28 that inhibited fewer kinases within this targeted enzymatic panel. For TTBK1/2 actives, defined as exhibiting ≤50 PoC for one of both kinases, we observed a consistent trend that, like parent AMG28, all analogs were more potent inhibitors of TTBK2 versus TTBK1. When the terminal alkyne substituent was unmodified, TTBK1/2 were most tolerant of the seven-membered (AMG28) and ring-opened analogs (**3**) when compared to analogs bearing six- and eight-membered rings (**1** and **2**, respectively). Capping the free hydroxy of AMG28 with a methyl group (**11**) was not favored, suggesting that the hydrogen bond donating functionality may be important for potency. Analogs of AMG28 that removed a methyl group from the center bearing the alcohol (**4** and **5**) did not result in improvements in activity. In contrast, those that increased steric bulk around that same carbon center by incorporating one or more ethyl group in place of a methyl resulted in more potent inhibitors of TTBK1/2. The loss in potency for the remaining three analogs (**6**–**8**) suggests that the pocket is not tolerant of homologation of the alkynyl substituent. The seven-membered analogs confirmed that the binding pockets of TTBK1 and TTBK2 are sensitive to even subtle changes in substitution around the hydroxy center (methyl to ethyl or to hydrogen) and do not accommodate deeper projection of this group into the back pocket (Figure 2).

### Kinome-Wide Selectivity Profiling Reveals that TTBK Lead Molecules Bind with High Affinity to a Small Percentage of Kinases

Encouraged by the promising inhibition profiles in the 17-membered kinase panel, AMG28 and the four TTBK actives (≤50 PoC for one or both TTBK1 and TTBK2) were next analyzed for kinome-wide selectivity. This screening was done at 1 μM via the DiscoverX *scan*MAX platform, which includes 403 wild-type (WT) human kinases but does not include TTBK1 or TTBK2. The kinase for which AMG28 was originally designed, NIK, is among the 403 WT kinases (Li *et al*., 2013). There is substantial overlap between the MRC and DiscoverX *scan*MAX kinase libraries, which allows the comparison of results on the same kinases in two different assay formats (enzymatic versus binding) at the same concentration. Selectivity results are included in two formats within Table 1. There is a column corresponding to the selectivity score (S_10_(1 μM)), which was calculated for each analog and is equal to the percentage of 403 WT human kinases that exhibit binding with a PoC <10 at 1 μM. A final column converts this selectivity score into the number of kinases that bound with a PoC <10 in the *scan*MAX panel. The specific WT kinases that bound with a PoC <10 at 1 μM in the *scan*MAX panel for the five compounds (AMG28, **1, 3, 9**, and **10**) are included in Figure S1.

This more comprehensive selectivity analysis revealed interesting selectivity trends. The ring-opened analog **3** demonstrated the poorest kinome-wide selectivity, suggesting that its increased conformational flexibility allows binding to a broader range of kinases. The six- and seven-membered ring containing analogs (**1** and AMG28, respectively) demonstrated improved kinome-wide selectivity compared to **3**. The ring systems maintain planarity of the scaffold and, based on the co-crystal structures, direct the alkyne into the back pocket (Figure 2). The modest selectivity of these compounds, however, still needs to be improved to provide a high-quality chemical probe. As the targeted enzymatic assay screening results suggested, each replacement of a methyl group with an ethyl group in **9** and **10** resulted in increased selectivity. Analog **9** demonstrated the most favorable kinome-wide selectivity of the TTBK actives, while **10** was less selective and mimicked the selectivity exhibited by parent compound AMG28. Analog **10** bears a very similar substituted alkyne to that of the most selective TTBK inhibitors published by Biogen (BGN18 and BGN31, Figure 1A), only differing by the choice to isolate a single enantiomer (S) versus our racemate (**10**).

### Orthogonal Enzymatic Studies Validate Binding and Inhibition of TTBK1/2 and Other Kinases

We then tested compounds in the TTBK1 and TTBK2 radiometric enzyme assays in dose–response format at Eurofins. Except for AMG28 which was equipotent on both kinases, all analogs were more potent inhibitors of TTBK2 than TTBK1. The analog containing a six-membered ring (**1**) was the weakest in the TTBK1/2 enzymatic assays and exhibited IC_50_ values >1 μM. While the ring-opened analog (**3**) and AMG28 were equipotent on TTBK1, differences in inhibition of TTBK2 surfaced. Both **3** and AMG28 demonstrated TTBK1 IC_50_ values in the 805–816 nM range, but **3** exhibited a TTBK2 IC_50_ = 384 nM and AMG28 exhibited a TTBK2 IC_50_ = 988 nM (Table 1). Interestingly, Biogen scientists also prepared **3** and reported a five-fold or greater loss in potency relative to AMG28 in their biochemical and cell-based assays (Halkina *et al*., 2021). When considering the enzymatic data and kinome-wide selectivity for **1**, AMG28, and **3**, seven-membered AMG28 is the best tool of these three due to its more optimal potency and selectivity. Moving to other analogs containing a seven-membered ring, we find that **9** and **10** are the most potent inhibitors of both TTBK1 and TTBK2 from the series (IC_50_ values <600 nM for TTBK1 and <300 nM for TTBK2, Table 1). This enzymatic result corroborates the single-concentration data, which predicted these two analogs to be the most potent inhibitors. It also aligns with the library prepared by Biogen, where they found the addition of an ethyl group provides a boost in potency due to displacement of a water molecule from the hydrophobic back pocket and enables van der Waals interactions between the ethyl group and lipophilic residues L81 and F177 (Halkina *et al*., 2021). When considering the enzymatic data and kinome-wide selectivity for **9** and **10, 9** demonstrates both the best enzymatic potency and kinome-wide selectivity and was selected as the most promising TTBK lead inhibitor from our series.

Given its selectivity and TTBK potency, we looked carefully into the potential off-targets of compound **9** that we had identified in the kinome-wide and 17-membered kinase panel selectivity experiments. We selected 19 total kinases from the S_35_(1 μM) fraction (Bosc et al., 2017) of the *scan*MAX kinase binding results and/or kinases potently inhibited in our selective enzymatic panel when tested at 1 μM (Table S1). All WT human kinases for which there was an available enzymatic assay kinase at Eurofins, RBC, or SignalChem (annotated in Table S2) within the S_35_(1 μM) fraction were tested in dose–response format. This only excluded PIP4K2C, for which a corresponding enzymatic assay was not available. Evaluation of **9** in the PIP4K2C NanoBRET cellular target engagement assay revealed it to bind this kinase with an IC_50_ = 576 nM (Table S2). We also added TAOK2, TTBK1, and TTBK2 for enzymatic follow-up because **9** was more potent on TAOK2 than TAOK1 in our enzymatic panel (Table S1), and TTBK1 and TTBK2 are not included in the *scan*MAX panel. Examination of this data, captured in Table S2, reveals that, in addition to TTBK1 and TTBK2, six kinases are potently inhibited by **9** with IC_50_ values ≤400 nM, and four of these kinases (MAP4K5, MYLK4, PIKfyve, and YSK4) have IC_50_ values ≤100 nM in the respective enzyme assays.

### Structural Studies Confirm that Indolyl Pyrimidinamines Assume a Canonical ATP-Competitive Binding Mode and Access the Back Pocket of TTBK1

Motivated to understand how these potent inhibitors bind to TTBK, we solved TTBK1 co-crystal structures with the four most potent TTBK inhibitors from our library (AMG28, **3, 9**, and **10**). As shown in Figure 2 and in agreement with the Biogen structures, our analogs target the back pocket via displacement of the catalytic lysine to access a narrow channel formed by M107, K63, L81, and F177 from the DFG motif that adopts an ‘in’ conformation. The solvent-exposed seven-membered ring shared by **9** and **10** makes these analogs very planar, which was similarly observed with **3**, and helps orient the substituent on the alkyne to interact with E77 and F177 in the back pocket. The aminopyrimidine ring makes two key hydrogen bonds with TTBK1 backbone hinge residues Q108 and Q110.

Interestingly, our structures also demonstrate that F45 at the tip of the P-loop is highly flexible and can adopt both ‘in’ and ‘out’ conformations. Ring-opened analog **3** induces an ‘in’ conformation of P-loop F45 (Figure 2B), whereas those that are locked into a ring (AMG28, **9**, and **10**) bind with F45 in an ‘out’ conformation (Figure 2A). We postulate that the inward movement of F45 stabilizes **3**, which is supported by a thermal shift value that is very similar to that of its ring-closed parent, AMG28 (Figure 2C). Importantly, these structural studies also informed design of a suitable tracer for enabling cellular target engagement assays.

The introduction of an ethyl group in **9** and **10** enabled space filling within the hydrophobic back pocket formed by the gatekeeper M107, the αC L81, and the DFG ‘in’ F177 that could not be achieved by AMG28. This enhanced binding mode is accompanied by an increase in kinase stabilization, which registers as a melting temperature (Tm) shift that is ∼1.2–1.8 °C higher than that observed with AMG28 (Figure 2C). Overall, the trend in thermal shifts was consistent with what we observed in enzymatic assays, with the most potent enzymatic inhibitors resulting in the largest thermal shifts (Figure 2C).

### Development of TTBK1 and TTBK2 Cellular Target Engagement Assays

To enable a cellular target engagement assay for both TTBK1 and TTBK2, we looked to the NanoBRET technology. While cDNA corresponding to the N- and C-terminally NanoLuciferase (NLuc) tagged versions of TTBK1 and TTBK2 were generously provided to our lab by Promega, a screen of the commercially available NanoBRET tracers from Promega demonstrated that none provided an adequate assay window suitable for target engagement assays. For our purposes, a NanoBRET tracer is defined as an ATP-competitive small molecule ligand validated to bind the kinase target of interest and modified to bear a red-shifted fluorophore. The NLuc-tagged kinase and NanoBRET tracer produce bioluminescence resonance energy transfer (BRET) when the tracer is bound to the ATP site, and this BRET can be competed away in a dose-dependent manner by ATP-competitive ligands. An IC_50_ value is then calculated and corresponds with the engagement of the kinase by a test compound in live cells (Vasta et al., 2018).

To enable a NanoBRET assay, we designed a fit-for-purpose tracer based on analog **3**. This parent was selected based upon its ease of synthesis from a commercially available indole (Figure S3) and its broader inhibition profile when screened kinome-wide (S_10_(1 μM) = 0.084), suggesting that it may be a useful tracer to enable NanoBRET assays for other understudied kinases. Furthermore, compound **3** is not locked into a specific orientation as part of a larger ring system, which we hypothesize would allow it to rotate sufficiently to adopt a favorable conformation once structurally modified. Based on our co-crystal structure of analog **3** bound to TTBK1 (Figure 2B), the indole nitrogen is solvent exposed. We opted for N-alkylation at this position to install our linker-appended fluorophore (Figure 3A). A short alkyl linker was chosen based on preliminary experiments with other kinases that suggested a propyl chain increases cellular permeability of resultant tracers versus those bearing short and long polyethylene glycol linkers.

**Figure 3.**
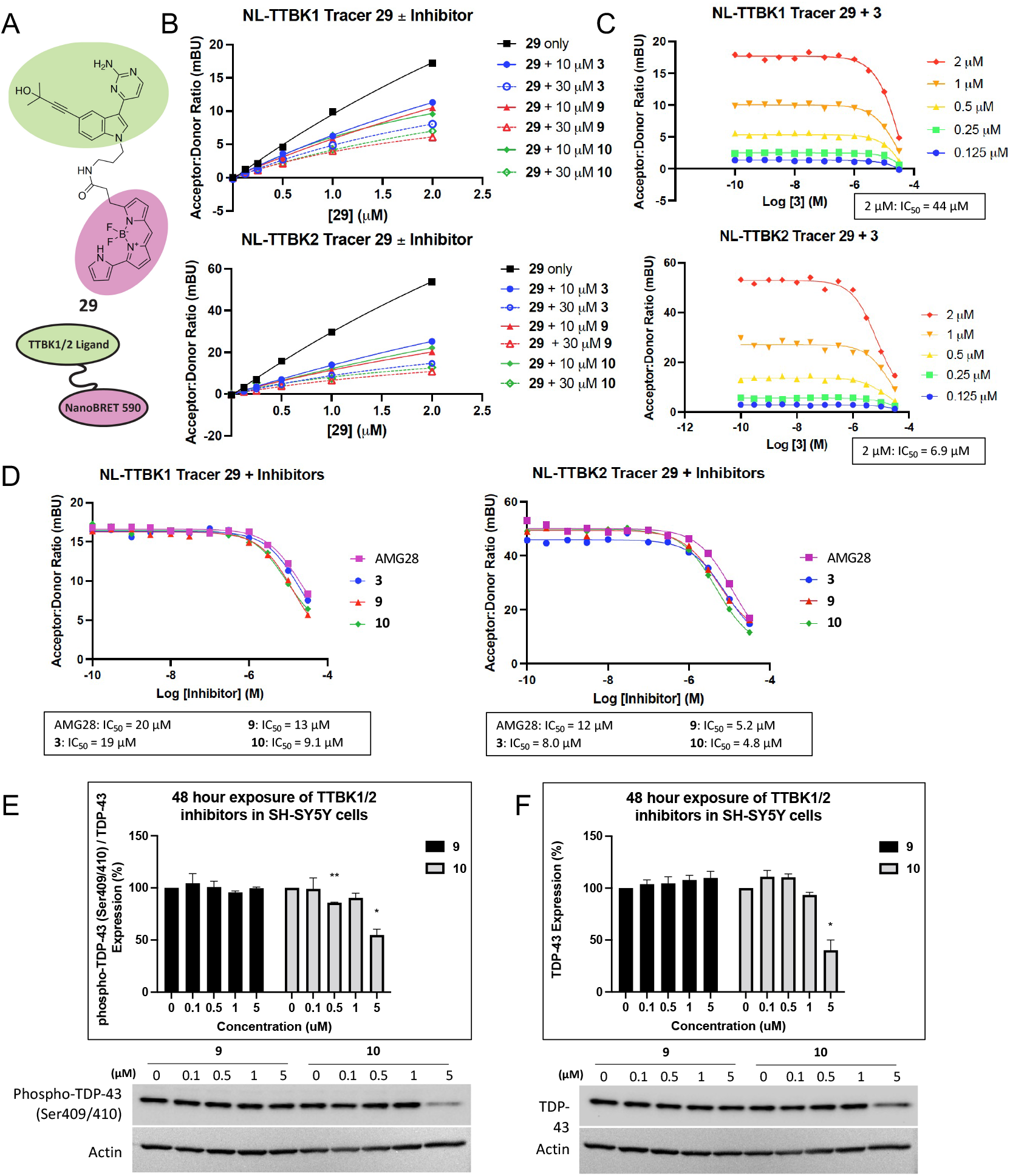
Cell-based studies of TTBK lead inhibitors. (A) Structure of tracer **29**. (B) Competition experiments between tracer **29** and analog **3, 9**, or **10** at either 10 or 30 μM for NLuc-TTBK1 and NLuc-TTBK2, n = 1. (C) Tracer titration results for NLuc-TTBK1 and NLuc-TTBK2 in the presence of analog **3**, n = 1. Calculated IC_50_ values are shown for each kinase at the recommended working tracer concentration of 2 μM. (D) Representative NanoBRET assay curves and corresponding IC_50_ values for active compounds AMG28, **3, 9**, and **10** in intact HEK293 cells, n = 1. (F) Western blot analyses of SH-SY5Y cells after 48 h treatment with **9** or **10**. Quantification of phospho-TDP-43 (Ser409/410, normalized to total TDP-43), n = 3. *p-value = 0.0011 and **p-value = 0.0154. (G) Western blot analyses of SH-SY5Y cells after 48 h treatment with **9** or **10**. Quantification of total TDP-43, n = 3. *p-value = 0.0260. For NanoBRET data and Western blots, error bars represent standard error of the mean (SEM). For Western blots, p-values were generated using a parametric unpaired t-test with Welch’s correction comparing each treatment condition to the untreated control.

Initial experiments aimed to determine which NLuc-tagged variant of each kinase (TTBK1 and TTBK2) yielded the best BRET signal and whether the tracer was able to permeate cells. Permeation studies were carried out by treating intact HEK293 cells and digitonin permeabilized HEK293 cells (both transiently transfected with the NLuc-TTBK1 and TTBK2 constructs) in dose–response format with 0.5–2 μM of tracer **29** (Figure S4). The N-terminally tagged version of both TTBK1 and TTBK2 yielded an optimal signal in the corresponding assays. Gratifyingly, a dose-dependent response of the BRET signal was observed in intact and permeabilized cells when treated with tracer **29**, confirming that the tracer enters HEK293 cells.

Before evaluating our TTBK inhibitors, we explored whether tracer **29** could be displaced from TTBK1 and TTBK2 by its unconjugated parent inhibitor (**3**). This competition experiment was used to confirm that the tracer binds in the same pocket as the parent compound instead of non-specifically or to an allosteric site. As shown in Figure 3B, for both TTBK1 and TTBK2, the BRET signal was decreased in a dose-dependent manner in the presence of different concentrations of tracer **29** through addition of 10 or 30 μM of parent **3**. Furthermore, a competition experiment using 10 or 30 μM of **9** or **10** also displaced tracer **29** (Figure 3B). The trend was maintained for analogs **3, 9**, and **10** when dosed at 10 and 30 μM, suggesting that **9** binds more efficaciously than **10**, and both have higher affinity than **3**.

After completing the competition experiments, we executed tracer titration experiments to determine the optimal working concentration of tracer **29** to use in the assays with NLuc-TTBK1 and NLuc-TTBK2. As shown in Figure 3C, parent compound **3** was dosed in the presence of increasing concentration of tracer: 0.125–2 μM. For both kinases, we found that 2 μM of tracer **29** afforded the best acceptor:donor ratio and provided the most optimal assay window for the respective NanoBRET assays. The EC_50_ of tracer **29** at its effective working concentration based on these tracer titration experiments is between 6–45 μM. The sub-optimal cellular penetrance of the tracer (Figure S4) is suggested to contribute to the observed EC_50_ values.

### Confirmation that Indolyl Pyrimidinamines Engage TTBK1 and TTBK2 in Live Cells

Preparation of tracer **29**, confirmation of its binding in cells and displacement, and determination of the optimal tracer concentration and preferred NLuc-tagged orientation for each kinase enabled NanoBRET assays for TTBK1 and TTBK2. Profiling the TTBK inhibitors was next executed in dose–response format. Results were obtained in intact as well as permeabilized HEK293 cells (Table 2). IC_50_ values for the majority of TTBK inhibitors were >30 μM in the live cell NanoBRET assay, except for AMG28, **3, 9**, and **10**, which were active in both assays. Two additional analogs (**1** and **5**) demonstrated IC_50_ values <30 μM in the TTBK2 NanoBRET assay only. AMG28, **3, 9**, and **10** exhibited IC_50_ values of 9–20 μM in the TTBK1 NanoBRET assay and 4–12 μM in the TTBK2 NanoBRET assay. In all cases, these analogs demonstrated more efficacious engagement of TTBK2 than TTBK1 in cells. This follows the trends in Table 1, where the same TTBK actives more potently inhibited TTBK2 than TTBK1. TTBK1 enzymatic IC_50_ values of 380–820 nM and TTBK2 enzymatic IC_50_ values of 175–990 nM translated to micromolar values in the respective NanoBRET assays for AMG28, **3, 9**, and **10**. This loss in potency in moving to cells is not unanticipated and indicates that additional optimization is warranted to improve both enzymatic and cellular potency of the leads. Our NanoBRET results in intact cells suggest that treatment with 5 μM of the active compounds would result in a phenotypic response in cell-based assays.

**Table 2.**
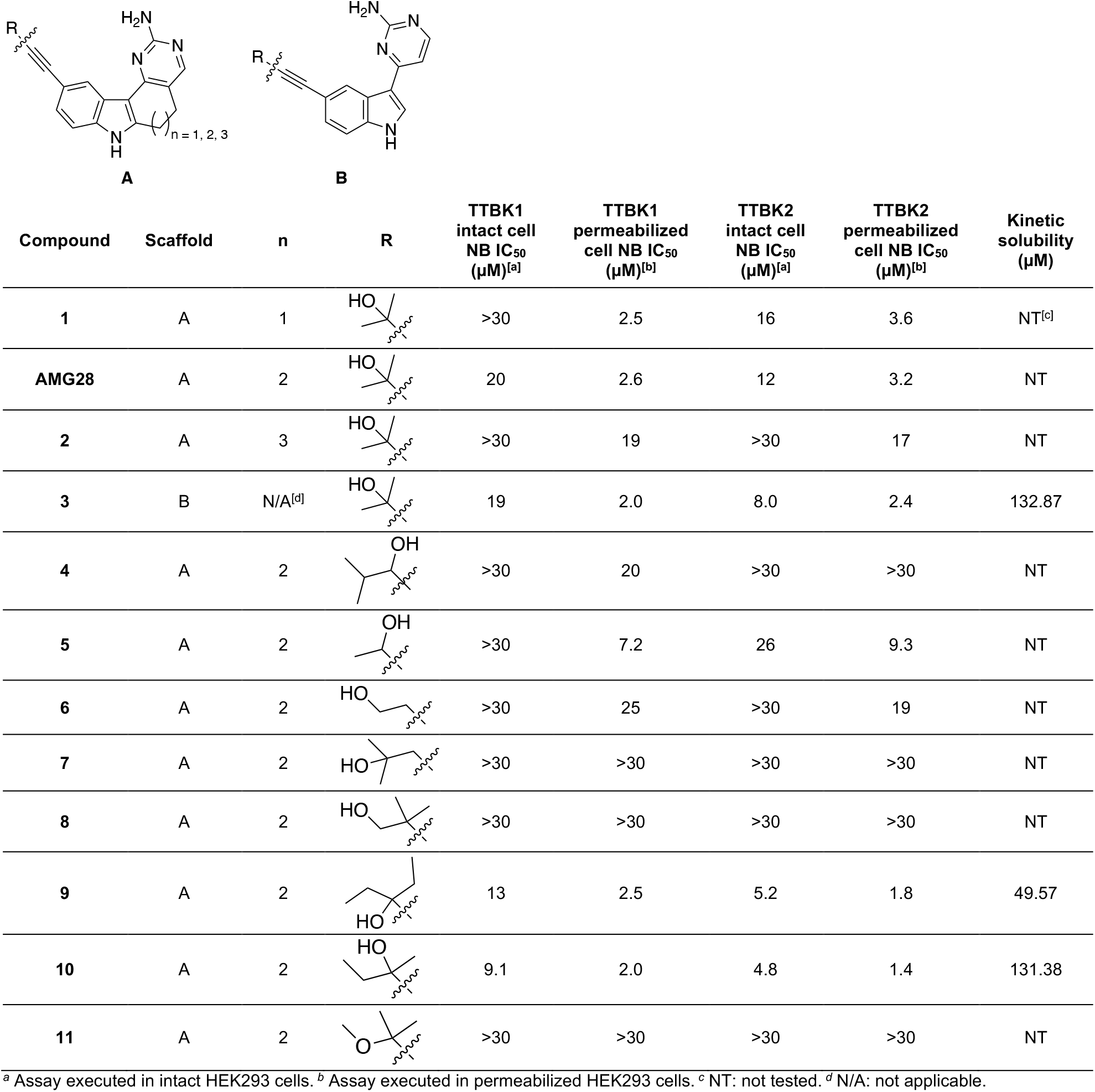
NanoBRET and Solubility Data for Indolyl Pyrimidinamine Library.

| Compound | Scaffold | n | R | TTBK1<br>intact cell<br>NB IC<sub>50</sub><br>(μM)<sup>[a]</sup> | TTBK1<br>permeabilized<br>cell NB IC<sub>50</sub><br>(μM)<sup>[b]</sup> | TTBK2<br>intact cell<br>NB IC<sub>50</sub><br>(μM)<sup>[a]</sup> | TTBK2<br>permeabilized<br>cell NB IC<sub>50</sub><br>(μM)<sup>[b]</sup> | Kinetic<br>solubility<br>(μM) |
| --- | --- | --- | --- | --- | --- | --- | --- | --- |
| 1 | A | 1 |  | >30 | 2.5 | 16 | 3.6 | NT<sup>[c]</sup> |
| AMG28 | A | 2 |  | 20 | 2.6 | 12 | 3.2 | NT |
| 2 | A | 3 |  | >30 | 19 | >30 | 17 | NT |
| 3 | B | N/A<sup>[d]</sup> |  | 19 | 2.0 | 8.0 | 2.4 | 132.87 |
| 4 | A | 2 |  | >30 | 20 | >30 | >30 | NT |
| 5 | A | 2 |  | >30 | 7.2 | 26 | 9.3 | NT |
| 6 | A | 2 |  | >30 | 25 | >30 | 19 | NT |
| 7 | A | 2 |  | >30 | >30 | >30 | >30 | NT |
| 8 | A | 2 |  | >30 | >30 | >30 | >30 | NT |
| 9 | A | 2 |  | 13 | 2.5 | 5.2 | 1.8 | 49.57 |
| 10 | A | 2 |  | 9.1 | 2.0 | 4.8 | 1.4 | 131.38 |
| 11 | A | 2 |  | >30 | >30 | >30 | >30 | NT |

<sup>a</sup> Assay executed in intact HEK293 cells. <sup>b</sup> Assay executed in permeabilized HEK293 cells. <sup>c</sup> NT: not tested. <sup>d</sup> N/A: not applicable.

When moving to permeabilized cells, AMG28, **3, 9**, and **10** exhibited IC_50_ values <3.2 μM in both NanoBRET assays, and IC_50_ values for the weaker TTBK inhibitors (**1, 2, 4, 5**, and **6**) improved as well (Table 2). Nine of twelve analogs demonstrated IC_50_ values <30 μM in the TTBK1 NanoBRET assay and eight of twelve analogs had IC_50_ values <30 μM in the TTBK2 NanoBRET assay. For both kinases, permeabilizing the cells improved the calculated IC_50_ values compared to those obtained in intact cells. Furthermore, the TTBK2 bias of the compounds was no longer observed when cells were permeabilized. Curves for the active compounds are included in Figure 3D (intact cells) and Figure S4 (permeabilized cells). The left-shifted curves in permeabilized cells suggests an opportunity to improve the cellular penetrance of the analogs, like the tracer, in future optimization efforts. To explore whether poor aqueous solubility was contributing to this result, **3, 9**, and **10** were sent for kinetic solubility measurements. Analog **9** was found to be less soluble than **3** and **10**, but all were in an acceptable range (Table 2).

### Analog 10 Modulates TTBK Downstream Signaling

To determine whether our inhibitors impact downstream signaling of TTBK1/2, we analyzed the ability of **9** and **10** to inhibit phosphorylation of TDP-43. TTBK1/2 belong to a group of only a few kinases that have been described to phosphorylate TDP-43 (Liachko *et al*., 2014; Nozal *et al*., 2022). We first explored the time-dependent effect of our compounds on TDP-43 phosphorylation in SH-SY5Y neuroblastoma cells. After 24 h of continuous treatment, there were no changes in phospho-TDP-43 or (total) TDP-43 protein expression when treated with up to 5 μM of **9**, and only (total) TDP-43 expression decreased (∼40%) at 5 μM with **10** (Figure S2). After 48 h of continuous treatment (Figure 3E and 3F), **9** did not change phospho-TDP-43 or total TDP-43 expression when treated with up to 5 μM. In contrast, 5 μM of **10** decreased phospho-TDP-43 expression by ∼45% and decreased total TDP-43 expression by ∼60%. Following continuous treatment for 72 h, 5 μM of **9** slightly decreased (∼13–16%) phospho- and total TDP-43 expression (data not shown). The concentration at which analog **10** reduced TDP-43 phosphorylation (5 μM) aligns well with its IC_50_ value in the TTBK1/2 NanoBRET assays (4.8–9.1 μM, Table 2).

Reports have suggested that exposure of SH-SY5Y cells to ethacrynic acid (EA) induces TDP-43 phosphorylation by causing cell death via glutathione depletion (Iguchi et al., 2012; Nozal *et al*., 2022). Following a published protocol, we treated SH-SY5Y cells with 5 μM of either **9** or **10** one hour prior to the addition of 40 μM EA (Nozal *et al*., 2022). Cells were harvested after 24 hours. We did not observe an enhancement of TDP-43 phosphorylation with EA treatment (Figure S2). One potential explanation is that our phospho-TDP-43 signal at baseline is rather robust in control cells. Of note, total TDP-43 was still decreased (∼44%) by 5 μM of **10**, but phospho-TDP-43 was not affected. This result was consistent with the 24 h continuous treatment without EA addition (Figure S2). In all experiments, limited toxicity was observed after 24 h continuous treatment with 5 μM of **9** or **10**. Based on the appearance of the cells and experiments carried out in parallel using our PIKfyve probe, we propose that at 5 μM these compounds are potently inhibiting PIKfyve (Drewry et al., 2022b). As shown in Table S2 and Figure S1, both compounds bind with high affinity to PIKfyve. PIKfyve inhibition leads to disruption of lysosome function that drives cell death (Gayle et al., 2017). LysoTracker staining overlaps with the enlarged vesicles observed at the 5 μM dose (Figure S2), which is consistent with PIKfyve inhibitor treatment (Drewry *et al*., 2022b; Gayle *et al*., 2017).

### Inhibition or Genetic Knockout of TTBK2 Reduce or Eliminate Cilia from Human Stem Cells

Due to their potency in binding and enzymatic assays and our interest in studying their impact on cilia, human induced pluripotent stem cells (iPSCs) were treated with either **9** or **10** and cilia were visualized. These cells were subjected to two different treatment protocols: either 72 h continuous starvation in the presence of inhibitor or 72 h continuous starvation with the inhibitor only present for the last 24 h. The impact of concentration as well as time course of treatment was evaluated. An example of the visual appearance of cells after 72 h continuous starvation in the presence of 1 μM of inhibitor is included in Figure 4A. Quantification of the percentage of ciliated cells in these single fields is included in Figure 4B. The percentage of ciliated cells over six visual fields was next calculated for each concentration and timepoint. As shown in Figures 4C and 4D, a time- and dose-dependent response of cilia to treatment with both compounds was observed but was more pronounced for **10**. Of the three concentrations and two timepoints examined, the most robust result was noted following 72 h continuous treatment and at a concentration of 1 μM with either analog.

**Figure 4.**
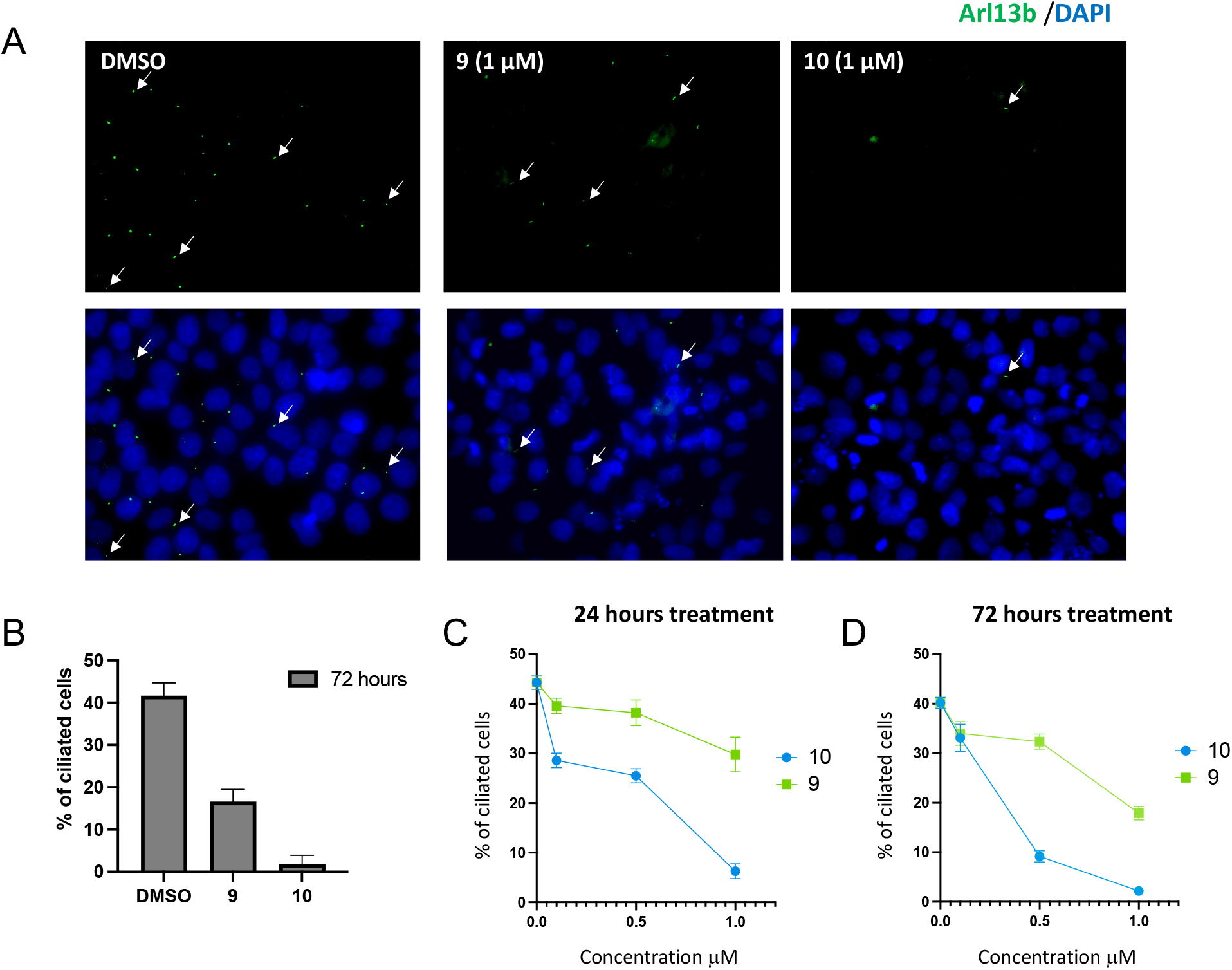
Images and quantification of cilia following treatment of human iPSCs with analog 9 or 10. (A) Visualization of cilia in human iPSCs following 72 h continuous starvation and treatment with DMSO or 1 μM of either **9** or **10** at 40x magnification. Arrows are used to highlight the location of some cilia in each panel and a legend of the stains used to visualize cilia and nuclei is placed above the panels. Primary cilia are labeled with the Arl13B antibody (green) and the nuclei with DAPI (blue). (B) Quantification of the percentage of ciliated cells per single field in (A). Error bars represent the SEM. (C,D) Quantification of the percentage of ciliated cells averaged from six random visual fields following 24 h (C) or 72 h (D) of continuous treatment plus starvation and in the presence of different concentrations of **9** or **10**. Error bars represent the SEM.

To compare with the results we observed when treating iPSCs with **9** and **10**, we generated TTBK2 KO iPSCs using the CRISPR/Cas9 system (Figure S5). We used the two guide RNA system to create a genomic deletion in the TTBK2 gene from exon 3 to exon 9. Two clones were isolated from the single cell screening. Sanger sequencing confirmed the genomic deletion from exon 3 to exon 9 on one allele and indels in exon 9 on the second allele of clone C26. The second clone, C15, displayed a deletion from exon 3 to exon 9 in one allele, but no indels in the second allele. Edited iPSCs remain pluripotent and retained their trilineage differentiation capabilities when compared to the isogenic cells (Figure S6). Figure 5A shows representative images of cells from the isogenic control cells, TTBK2 edited cells (C15 and C26), and an unrelated human iPSC line. Examination of these images clearly shows that cilia are absent in TTBK2 KO C26 and modestly reduced in TTBK2 KO C15. The lack of cilia expression in TTBK2 KO C26 was observed even in the absence of starvation (Figure S6). These findings are consistent with what was previously reported with respect to cilia in the brains of conditional TTBK2 KO adult mice (Bowie and Goetz, 2020). We quantified the percentage of ciliated cells when ten fields were visualized and averaged in these various iPSC lines and plotted those averages versus what was observed after 72 h starvation and continuous treatment with **10**. This graph in Figure 5B demonstrates that treatment of iPSCs with **10** phenocopies the TTBK2 KO C26 cells.

**Figure 5.**
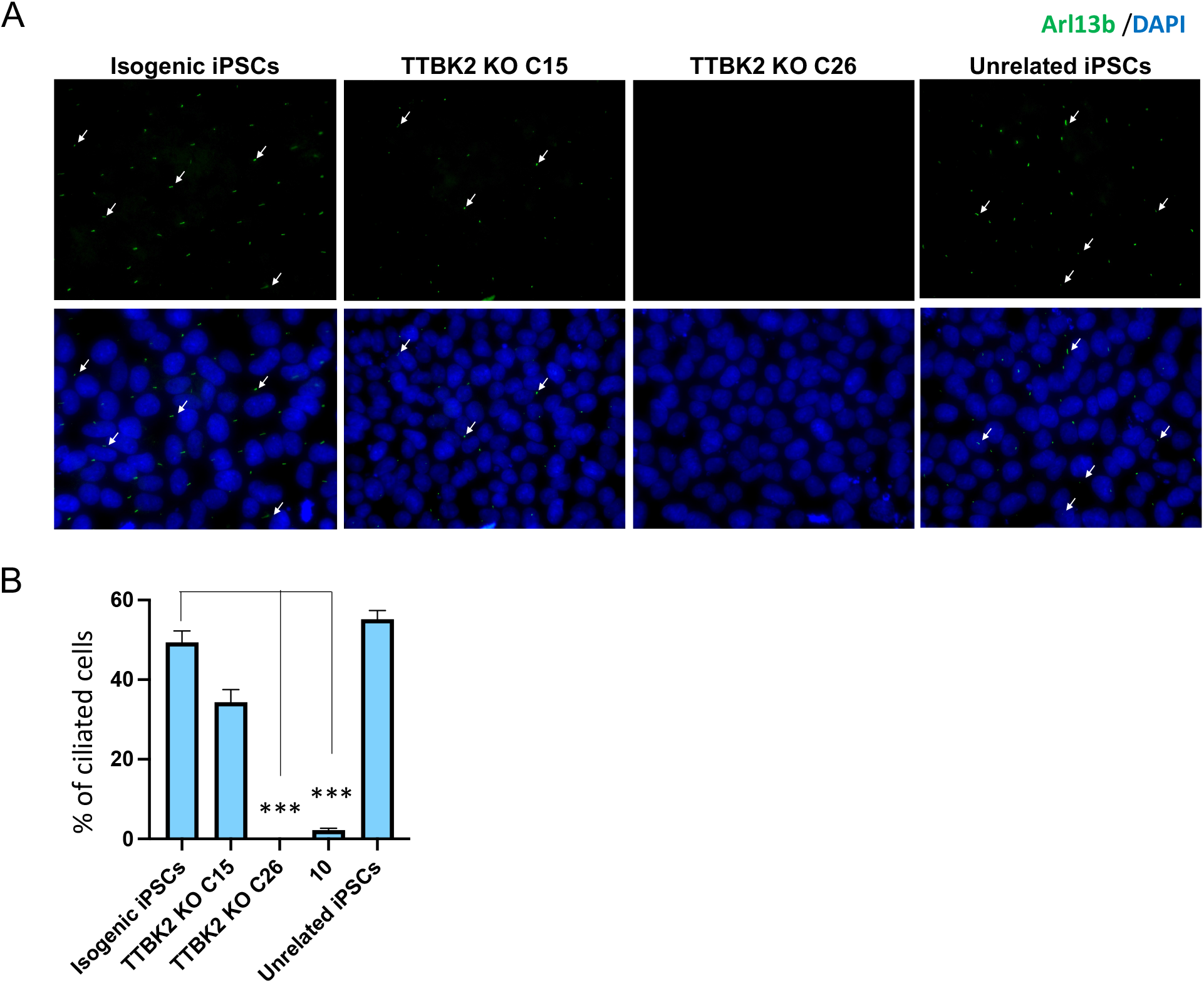
Images and quantification of cilia in human iPSCs compared to TTBK2 KO iPSCs. (A) Visualization of cilia in human isogenic iPSCs, TTBK2 KO C15, TTBK2 KO C26, and unrelated iPSCs following 72 h starvation. Arrows are used to highlight the location of some cilia in each panel and a legend of the stains used to visualize cilia and nuclei is placed above the panels. Primary cilia are labeled with the Arl13B antibody (green) and the nuclei with DAPI (blue). (B) Plot of the percentage of cells expressing cilia averaged from ten random visual fields for human isogenic iPSCs, TTBK2 KO C15, TTBK2 KO C26, and unrelated iPSCs versus percentage of cells expressing cilia averaged from six random visual fields following 72 h of continuous treatment with 1 μM of **10**. Error bars represent the SEM and p-values calculated by comparing the percentage of cells expressing cilia from three different biological experiments using an unpaired Student’s t-test. ***p-value = 0.0001.

## DISCUSSION

While we hypothesized that minor structural changes to the alkyne of AMG28 would result in the production of analogs of similar potency and selectivity, compounds with more significant structural changes, such as changes in ring size or ring opening, were designed to probe the impact of specific structural modifications on potency and selectivity for TTBK1 and TTBK2. The orthogonal data collected for various analogs, together with the co-crystal structures, allows the development of structure–activity relationships (SAR) for the indolyl pyrimidinamine scaffold. Gratifyingly, a slight bias of our compounds to bind to and inhibit TTBK2 more efficaciously than TTBK1 was observed across the biochemical and cell-based analysis methods. Furthermore, the potency of the TTBK active compounds was maintained in these binding and enzymatic assay formats.

With both a six-membered (**1**) and eight-membered analog (**2**) of AMG28 in which only the size of the ring system was modified, TTBK1/2 potency was significantly altered. The enzymatic data for this small series suggests that the seven-membered ring offers enhanced efficacy versus the smaller and larger ring systems. Based on our co-crystal structures (Figure 2), the actual methylene units that make up these rings are solvent-exposed, and the systems are largely planar. Thus, it is likely that the ring system size alters the orientation and trajectory of the alkyne and the ability of the compounds to make key contacts with the back pocket is diminished with analog **1** and especially with analog **2** when compared to AMG28. When the ring system was removed to yield analog **3**, potency was maintained across enzymatic, binding, and cell-based assays when comparing AMG28 and analog **3** (Tables 1, 2, and S1). Importantly, the NanoBRET assay results (Table 2) indicate that AMG28 and analog **3** share similar cellular penetrance capacity and maintain TTBK kinase binding in cells. A notable difference between AMG28 and analog **3** is kinome-wide selectivity. Ring-opened analog **3** binds a larger portion of the screenable kinome than AMG28, which could be orchestrated by the enhanced rotational freedom of **3** that allows it to adopt alternative conformations within the ATP sites of different kinases.

More subtle structural changes are present when examining analogs **4**–**11**. Comparison of the data in Tables 1, 2, and S1 for analogs **9** and **10** to that generated for AMG28 confirms that the portion of the binding pocket that accommodates the alkyne is sensitive to changes in steric bulk. This back pocket, which is accessed via a narrow channel in TTBK1, is filled by hydrophobic groups on **9** and **10** that enhance their binding versus AMG28 (Figure 2A). It is also worth noting that all of our inhibitors, including AMG28, that protrude into the back pocket produce remarkably higher Tm shift values than TTBK inhibitors that are unable to access the back pocket, such as those reported recently by Nozal et al. built upon the AZ2 scaffold (Figure 1, ΔTm <3.5) (Nozal *et al*., 2022). These data suggest that space filling the TTBK1/2 back pocket is a design strategy via which to develop analogs of **9** and **10** with increased kinase stabilization and enhanced inhibitory potency. Examination of other analogs bearing a seven-membered ring (**4**–**8** and **11**), however, supports the idea that the placement of hydrophobic bulk must be very deliberate, and branching from the carbon directly attached to the alkyne with two carbon chains is optimal (AMG28, **9**, and **10** versus **4** and **5**). Coupled with this finding is the discovery that the hydroxyl group on the alkyne terminus cannot be moved from this carbon without losses in TTBK potency (AMG28, **9**, and **10** versus **6**–**8**). Finally, analog **11** demonstrates that the hydroxyl group cannot be capped with a methyl group without dramatic losses in activity, indicative of forfeiture of a key interaction and/or interference of the resultant methyl ether with the back pocket.

Comparison of the kinome-wide selectivity of lead analogs **9** and **10** suggests that while TTBK1 and TTBK2 prefer the addition of a single methylene unit in **9**, many other kinases do not tolerate this structural modification. As a result, increased potency of analog **9** versus **10** is accompanied by enhanced kinome-wide selectivity. Fifteen kinases fewer kinases are in the S_10_(1 μM) fraction when the methyl in analog **10** is homologated to an ethyl in analog **9**. Thus, filling the pocket with properly oriented hydrophobic bulk is a way to increase both potency for TTBK1 and TTBK2 and preclude the binding of off-target kinases. Our finding that the methyl, ethyl substitution on analog **10** resulted in a boost in TTBK potency was echoed by Biogen in their discovery of BGN18 and BGN31 (Figure 1), which bear an alkyne with the same substitution as **10** (Halkina *et al*., 2021). BGN18 and BGN31 were scanned in the same DiscoverX *scan*MAX panel at 1 μM and were found to bind with PoC <10 sixteen and eight kinases, respectively (Halkina *et al*., 2021). Many of these off-target kinases are bound within the S_10_(1 μM) fraction by our TTBK actives and were potently inhibited in the MRC panel (Figure 1B). This side-by-side comparison makes **9**, which binds seven kinases with PoC <10 at 1 μM (Table 1), the TTBK inhibitor with the lowest kinome-wide selectivity score from these structurally related series.

The modest enzymatic (150–400 nM, Table 1) and NanoBRET (5–13 μM, Table 2) IC_50_ values for TTBK1 and TTBK2 coupled with the potent inhibition of off-target kinases support that further optimization of **9** is required to furnish a TTBK chemical probe. Our cellular studies, examining target engagement, downstream phosphorylation, and phenotypic results, suggest that the cellular penetrance of these analogs is suboptimal. The planarity of our analogs (Figure 2) is a feature that we suggest could be modified to improve their solubility and may result in improved cellular penetrance. The loss in activity when moving from TTBK biochemical assays into a cellular system is not unique to these kinases or our inhibitors. Compound **9** exhibits nearly a 34-fold loss in activity when moving from the TTBK1 enzymatic to TTBK1 NanoBRET assay. This is directly in line with the Biogen analogs that demonstrate nearly 14- and 117-fold losses in activity (BGN18 and BGN31, respectively) when moving from a TTBK1 enzymatic assay to a cellular assay that measured tau phosphorylation on Ser422 in HEK293T cells (Halkina *et al*., 2021). Dosing with these compounds at micromolar concentrations >1 μM is non-ideal since we have not assessed their kinome-wide selectivity at higher doses (collected at 1 μM). Fortunately, dosing with analog **9** at sub-micromolar concentrations demonstrated a significant ciliogenesis phenotype in human iPSCs (Figure 4).

All currently employed cell-based assays used to validate and measure the activity of TTBK inhibitors in cells rely on an indirect measurement, such as tau or TDP-43 phosphorylation (Halkina *et al*., 2021; Nozal *et al*., 2022). There is no commercial or published assay that directly measures engagement of TTBK1/2 by a small molecule in cells. Our TTBK1/2 NanoBRET assays offer direct assessment of binding to TTBK1 or TTBK2 in live cells. Furthermore, these assays allow evaluation of binding kinetics, compound residence time, and cellular penetrance and can be used to drive iterative medicinal chemistry optimization (Robers et al., 2015).

The results we observed with respect to cilia are aligned with literature reports related to the roles of TTBK1/2 in ciliogenesis. Several labs have worked to define the mechanism via which TTBK2 modulates this process and the physiological substrates involved. Recruited by CEP164, TTBK2 has been found to colocalize with CEP83 at the root of the centriole distal appendage (Čajánek and Nigg, 2014; Lo et al., 2019; Sánchez and Dynlacht, 2016). Phosphorylation of CEP83 by TTBK2 is essential for the early ciliogenesis steps, including ciliary vesicle docking and CP110 removal from the mother centriole, which initiate assembly of distal appendages to form cilia (Lo *et al*., 2019; Sánchez and Dynlacht, 2016). Specifically, TTBK2, in concert with other proteins and phosphatidylinositol, has been suggested to engage in remodeling of the distal ends of mother centrioles and basal bodies to allow for activation of microtubule growth and axonemal extension (Sánchez and Dynlacht, 2016). Other essential ciliogenesis mediators, including CEP167 and CEP97, have been characterized as TTBK2 substrates as well (Čajánek and Nigg, 2014; Oda et al., 2014; Sánchez and Dynlacht, 2016). A lack of validated antibodies that bind to centriole proteins has made it difficult to distinguish the precise role of TTBK2 phosphorylation in these ciliogenesis-promoting processes.

Conditional KO of TTBK2 in adult mice results in degenerative cerebellar phenotypes. These mice quickly lose cilia throughout their brain (Bowie and Goetz, 2020). In addition, a TTBK2-null mouse mutant was found to lack primary cilia and exhibit defects in hedgehog signaling. Loss of TTBK2 in this mouse model abolished the recruitment of IFT proteins essential for anterograde and retrograde trafficking as well as axoneme extension and maintenance. These TTBK2-null mice had basal bodies that lacked axonemes (Goetz et al., 2012). These *in vivo* models align with our findings of cilia frequency reduction in TTBK2 KO iPSCs and upon treatment of human iPSCs with our TTBK inhibitors. The next step will be to identify an optimized TTBK2 chemical probe to further characterize the role of TTBK2 in mediating ciliogenesis and to explore how TTBK2 loss of function propagates diseases, including SCA11 and others in which cilia abnormalities are pathogenic.

## Supporting information

Combined Supplemental Data

## SIGNIFICANCE

**We have described the design, synthesis, and biological evaluation of a series of indolyl pyrimidinamines as chemical tools to interrogate the functions of TTBK1 and TTBK2. We have demonstrated that our analogs are inhibitors of TTBK1 and TTBK2 and exhibit a slight bias for TTBK2 using three orthogonal assay formats: radiometric enzyme assays at Eurofins, thermal shift assays, and NanoBRET cellular target engagement assays in live cells for both TTBK1 and TTBK2. No commercially available or published assay measures TTBK1 or TTBK2 engagement in cells, so the TTBK NanoBRET assays described here are a useful addition to the field. Solving co-crystal structures of lead analogs with TTBK1 provided insights that enabled the development of TTBK1 and TTBK2 NanoBRET assays. These structures illustrated that the potency of our analogs was derived in part from their ability to access the back pocket of the TTBK1 ATP binding site. Analog 10, which was confirmed to inhibit downstream TTBK signaling, emerged as a promising tool compound to assess the role of TTBK1/2 in ciliogenesis. This cell active inhibitor of TTBK2, which binds with high affinity to only a small subset of the screenable human kinome, reduced cilia frequency when dosed in human stem cells. As TTBK2 KO resulted in human stem cells that completely lacked cilia and analog 10 phenocopied this observation, compound 10 can be used as a tool molecule to help characterize TTBK2-mediated pathways in cells. These studies may be focused on building further understanding of the roles TTBK2 plays in ciliogenesis but could also help identify as yet unknown functions of TTBK2. The involvement of cilia in a diverse spectrum of diseases enhances the importance of tools that facilitate efforts to study and understand all aspects of cilia biology. Studies using analog 10 will help deconvolute the distinct roles of tau tubulin kinases and validate additional substrates of these understudied kinases in key disease-propagating pathways**.

## ACKNOWLEDGEMENTS

Constructs for NanoBRET measurements of TTBK1, TTBK2, and PIP4K2C were kindly provided by Promega. We used the TREE*spot* kinase interaction mapping software from DiscoverX to prepare the kinome trees included in Supplemental Information: http://treespot.discoverx.com. We thank ChemSpace LLC for synthetic support and the beamline scientists at the Swiss Light Source (CH) for their assistance with data collection. Input and expertise offered by Sarah Goetz was appreciated by the authors.

SGC, a registered charity (number 1097737) that receives funds from Bayer AG, Boehringer Ingelheim, the Canada Foundation for Innovation, Eshelman Institute for Innovation, Genentech, Genome Canada through Ontario Genomics Institute [OGI-196], EU/EFPIA/OICR/McGill/KTH/Diamond, Innovative Medicines Initiative 2 Joint Undertaking [EUbOPEN grant 875510], Janssen, Merck KGaA (aka EMD in Canada and USA), Pfizer, the São Paulo Research Foundation-FAPESP, and Takeda. Research reported in this publication was supported in part by the NC Biotech Center Institutional Support Grant 2018-IDG-1030, NIH U24DK116204, and DoD ALSRP award AL190107.

## Author Contributions

Conceptualization, F.M.P., D.H.D., A.C., S.H., A.S.B., and A.D.A.; Methodology, F.M.P., A.A.B., A.S.B., and A.D.A.; Validation, F.M.P., A.C., S.H., A.S.D., A.S.B., and A.D.A.; Formal Analysis, F.M.P., A.C., S.H., A.S.D., J.L.S., and A.S.B.; Investigation, F.M.P., A.B.M., A.C., S.H., A.A.B., and J.L.S.; Resources, F.M.P., A.C., S.H., J.L.S., A.S.B., and A.D.A.; Writing – Original Draft, A.D.A.; Writing – Review & Editing, all authors; Visualization, F.M.P., A.C., S.H., A.S.B., and A.D.A.; Supervision, A.C., A.S.B., and A.D.A.; Project Administration, A.D.A.; Funding Acquisition, A.D.A.

## STAR METHODS

### RESOURCE AVAILABILITY

#### Lead Contact

Further information and requests for resources and reagents should be directed to and will be fulfilled by the Lead Contact, Alison Axtman.

#### Materials Availability

All compounds can be requested through contacting the Lead Contact.

#### Data and Code Availability

This study did not generate/analyze datasets/code.

### EXPERIMENTAL MODEL AND SUBJECT DETAILS

#### Cell Lines

##### HEK293

Human embryonic kidney (HEK293) cells (hypotriploid, female, fetal) were obtained from ATCC and cultured in Dulbecco’s Modified Eagle’s medium (DMEM, Gibco) supplemented with 10% (v/v) fetal bovine serum (FBS, Corning). HEK293 cells were incubated in 5% CO_2_ at 37°C and passaged every 72 hours with trypsin, not allowing them to reach confluency.

##### SH-SY5Y

Human neuroblastoma (SH-SY5Y) cells were obtained from ATCC and cultured in DMEM/F12 medium (Gibco) supplemented with 10% (v/v) fetal bovine serum (FBS, Corning). SH-SY5Y cells were incubated in 5% CO_2_ at 37°C and passaged every 48 hours with trypsin, not allowing them to reach confluency.

##### Wild-type human iPSCs

Human Gibco (Thermo Fisher Scientific A18945) and UNCC001-A (Molina et al., 2020) iPSCs were cultured on Matrigel-coated dishes in StemFlex medium (Thermo Fisher Scientific A3349401). Cells were passaged twice a week with 0.5 mM EDTA at a 1:10 ratio. Cell media was supplemented with 10 μM Y27632 (Selleckchem S1049) for 24 hours after each passage.

##### TTBK2 KO iPSCs

The two guide RNA (gRNA) target approach was used to create a genomic deletion in the TTBK2 gene. Guide RNAs were designed to target the exon 3 (TGGAGAAATTTACGATGCCT) and exon 9 (GGAGCAAGTAGGCTCTATTA) simultaneously. The ribonucleoprotein (RNP) system was used as described previously (Battaglia et al., 2019). Briefly, 3 × 10^5^ Gibco iPSCs were electroporated with 900 ng of each single gRNA using the Neon Transfection System (Thermo Fisher Scientific). 72 hours after electroporation, iPSCs were dissociated into single cells and plated on Matrigel-coated 96-well plates. Single-cell colonies were selected after two weeks, and the genomic DNA was extracted and tested for gene deletion by PCR. Sanger sequencing of positive clones demonstrated gene deletion in one allele in clone 15 and two alleles for clone 26. TTBK2 lack of expression was confirmed by mRNA (using the primers Ex3-Fw-TCCGGCTGGGTAGACAGATT and Ex4-Rv-AGCGAAGTTCGACGGTTTGA) (Figure S5).

Stemness and trilineage differentiation capabilities of the TTBK2 KO clones was performed as previously described (Beltran et al., 2021) (Figure S6). TTBK2 KO iPSCs were cultured as described in the section above.

### METHOD DETAILS

#### General Information for Chemical Synthesis

Reagents were obtained from verified suppliers and employed without purification. Temperatures are in degrees Celsius (°C); solvent was removed using a rotary evaporator; and thin layer chromatography as well as LC–MS were used to monitor reaction progress. ^1^H NMR and additional analytical data was collected for intermediates and final compounds to verify their identity and evaluate their purity. ^1^H and ^13^C NMR spectra were obtained in DMSO-*d*_6_ or MeOD-*d*_4_ and recorded using Bruker spectrometers. Magnet strength is listed in each experimental write-up along with peak positions in parts per million (ppm). Peaks in NMR spectra are calibrated versus the shift of the indicated deuterated solvent and coupling constants (*J* values) are reported in hertz (Hz). Peak multiplicities are included as follows: singlet (s), doublet (d), doublet of doublets (dd), triplet (t), pentet (p), and multiplet (m). Purity of all compounds was assessed via HPLC.

Preparation of AMG28 as well as analogs **4**–**11** began with Fischer indole synthesis to produce **15**. Synthesis of analogs **1** and **2** was made possible from commercially available indoles **14** and **16**, respectively. Intermediates **17**–**19** were accessed from indoles **14**–**15** via selective oxidation with DDQ. Next, intermediates **20**–**22** were synthesized from **17**–**19** by refluxing with Bredereck’s reagent and then cyclized to the corresponding aminopyrimidines **23**–**25** using guanidine. The desired TTBK inhibitors **1, 2, 4**–**11**, and AMG28 were synthesized from **23**–**25** via Sonogashira coupling conditions (Figure S3). In analogous fashion, inhibitor **3** was accessed from commerically available indole **26** using Sonogashira coupling conditions (Figure S3). Full synthetic procedures and compound characterization are included in a separate manuscript (Drewry *et al*., 2022b). ^1^H NMR data and spectra for the cell active TTBK inhbitors are included below to confirm identity.

Preparation of Tracer **29** began with N-alkylation of the indole nitrogen of commerically available indole **26** with a Boc-protected propyl amine to produce **27**. Next, acidic conditions promoted the liberation of the pendant amine in intermediate **28**. A two-step sequence was employed to first install the desired alkyne bearing a propargylic alcohol via Sonogashira coupling conditions, and then attach the NanoBRET 590 dye under basic conditions and yield final tracer **29** (Figure S3). It is worth noting that the propargylic alcohol is acid labile and should not be exposed to acidic conditions, even during purification. Full characterization of the final tracer is included.

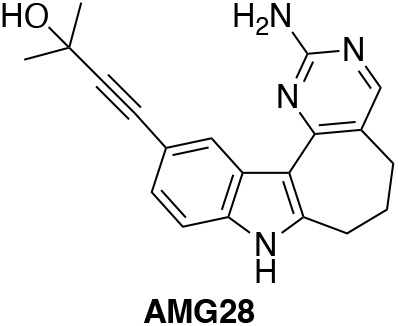

*4-(2-amino-5,6,7,8-tetrahydropyrimido[4’,5’:3,4]cyclohepta[1,2-b]indol-11-yl)-2-methylbut-3-yn-2-ol (AMG28)*. The analytical data for AMG28 matches that previously published (Drewry *et al*., 2022b; Li *et al*., 2013). ^1^H NMR (400 MHz, DMSO-*d*_*6*_) δ 11.61 (s, 1H), 8.71 (s, 1H), 8.29 (s, 1H), 7.92 (s, 1H), 7.26 (d, *J* = 8.3 Hz, 1H), 7.11 (dd, *J* = 8.4, 1.6 Hz, 1H), 6.20 (s, 2H), 3.19 – 3.15 (m, 2H), 2.66 – 2.59 (m, 2H), 1.98 – 1.92 (m, 2H), 1.50 (s, 6H).

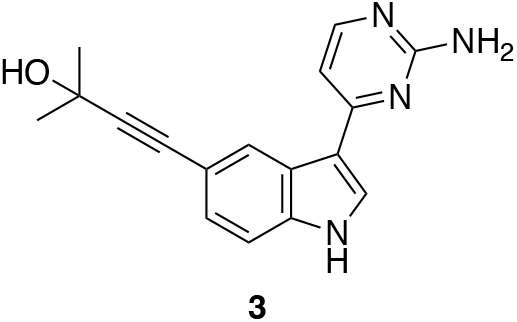

*4-(3-(2-aminopyrimidin-4-yl)-1H-indol-5-yl)-2-methylbut-3-yn-2-ol (****3***). The analytical data for **3** matches that previously published (Drewry *et al*., 2022b; Halkina *et al*., 2021). ^1^H NMR (400 MHz, DMSO-*d*_*6*_) δ 11.83 (s, 1H), 8.56 (s, 1H), 8.15 – 8.09 (m, 2H), 7.41 (d, *J* = 8.4 Hz, 1H), 7.17 (d, *J* = 8.9 Hz, 1H), 7.01 (d, *J* = 5.3 Hz, 1H), 6.50 (s, 2H), 1.50 (s, 6H).

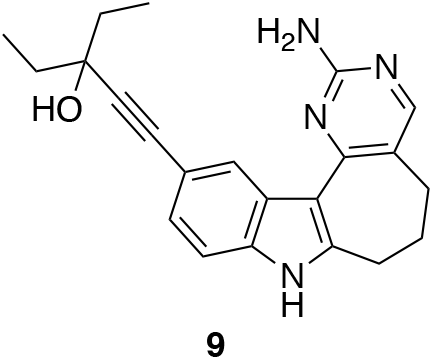

*1-(2-amino-5,6,7,8-tetrahydropyrimido[4’,5’:3,4]cyclohepta[1,2-b]indol-11-yl)-3-ethylpent-1-yn-3-ol (****9***). The analytical data for **9** matches that previously published (Drewry *et al*., 2022b). ^1^H NMR (400 MHz, DMSO-*d*_*6*_) δ 11.60 (s, 1H), 8.70 (d, *J* = 1.6 Hz, 1H), 7.27 (d, *J* = 8.3 Hz, 1H), 7.13 (dd, *J* = 8.2, 1.6 Hz, 1H), 6.14 (s, 2H), 5.02 (s, 1H), 3.17 (t, *J* = 6.5 Hz, 2H), 2.66 – 2.59 (m, 2H), 1.97 (dd, *J* = 10.5, 5.5 Hz, 2H), 1.75 – 1.56 (m, 4H), 1.03 (t, *J* = 7.4 Hz, 6H).

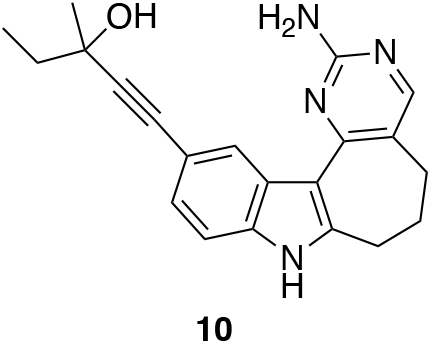

*1-(2-amino-5,6,7,8-tetrahydropyrimido[4’,5’:3,4]cyclohepta[1,2-b]indol-11-yl)-3-methylpent-1-yn-3-ol (****10***). The analytical data for **10** matches that previously published (Drewry *et al*., 2022b). ^1^H NMR (400 MHz, DMSO-*d*_*6*_) δ 11.60 (s, 1H), 8.71 (s, 1H), 8.16 (d, *J* = 2.2 Hz, 1H), 7.92 (s, 1H), 7.27 (d, *J* = 8.2 Hz, 1H), 7.12 (dd, *J* = 8.3, 1.7 Hz, 1H), 6.17 (s, 2H), 3.17 (t, *J* = 6.6 Hz, 2H), 2.62 (d, *J* = 9.1 Hz, 2H), 1.98 – 1.92 (m, 2H), 1.71 – 1.62 (m, 2H), 1.45 (s, 3H), 1.03 (t, *J* = 7.4 Hz, 3H).

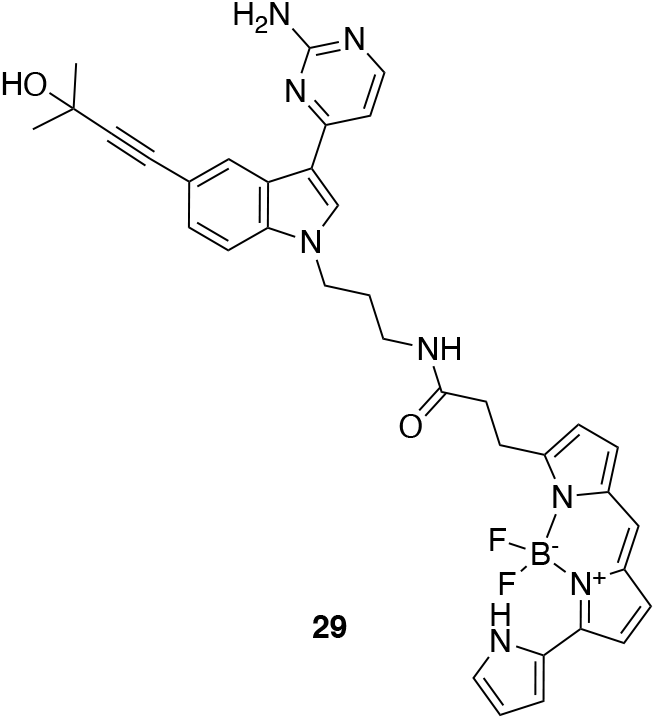

*N-(3-(3-(2-aminopyrimidin-4-yl)-5-(3-hydroxy-3-methylbut-1-yn-1-yl)-1H-indol-1-yl)propyl)-3-(5,5-difluoro-7-(1H-pyrrol-2-yl)-5H-5λ4,6λ4-dipyrrolo[1,2-c:2’,1’-f][1,3,2]diazaborinin-3-yl)propenamide (****29***). ^1^H NMR (400 MHz, MeOD-*d*_*4*_) δ 8.58 – 8.56 (m, 1H), 8.01 (d, *J* = 5.5 Hz, 1H), 7.99 (s), 7.37 (dd, *J* = 8.6, 0.8 Hz, 1H), 7.24 (dd, *J* = 8.5, 1.6 Hz, 1H), 7.21 – 7.17 (m, 2H), 7.14 (d, *J* = 4.6 Hz, 1H), 7.12 (s, 1H), 6.99 (d, *J* = 4.6 Hz, 1H), 6.95 (d, *J* = 5.5 Hz, 1H), 6.87 (d, *J* = 3.9 Hz, 1H), 6.36 – 6.33 (m, 2H), 4.14 (t, *J* = 7.0 Hz, 2H), 3.23 (t, *J* = 6.5 Hz, 2H), 2.67 (t, *J* = 7.5 Hz, 2H), 2.03 (p, *J* = 6.8 Hz, 2H), 1.60 (s, 6H). LCMS: calculated for C_36_H_36_BF_2_N_8_O_2_ [M + H]^+^: 661.30. Found: 661.34.

#### Enzymatic Assays

Eurofins kinase enzymatic radiometric assays were executed at the K_m_ = ATP using a single concentration (1 μM) in duplicate to generate PoC values for TTBK1 and TTBK2 in Table 1 as well for as each kinase listed in Table S1. Eurofins kinase enzymatic radiometric assays were executed at the K_m_ = ATP in dose– response (9-pt curve) format for TTBK1 and TTBK2, included in Table 1 as enzymatic IC_50_ values, as well as for the majority of kinases in Table S2. Details about the substrate used, protein constructs, controls, and assay protocol for these kinase assays are available at the Eurofins website: https://www.eurofinsdiscoveryservices.com. Reaction Biology Corp. (RBC) radiometric HotSpot kinase assays were carried out at the K_m_ = ATP in dose–response (10-pt curve) format for MYLK4, RIPK5, and YSK4 in Table S2. Details about the substrate used, protein constructs, controls, and assay protocol for these kinase assays are available at the RBC website: https://www.reactionbiology.com/list-kinase-targets-us-facility. SignalChem developed an ADP-Glo assay to test for enzymatic inhibition of PIKfyve in dose– response (10-pt curve) format in duplicate (Drewry *et al*., 2022b).

#### Kinome Screening

The Eurofins DiscoverX Corporation *scan*MAX assay platform was employed to assess the selectivity of specific analogs at a single concentration (1 μM). As described previously, this assay platform profiles a compound against 403 wild-type human kinases and delivers percent of control values (Davis et al., 2011). These percent of control values are included in Table 1 and corresponding kinome tree diagrams as well as tables for each analog included in Figure S1.

#### General Information for NanoBRET assays

HEK293 cells were transfected with constructs of TTBK1 or TTBK2 tagged with NLuc on the N- or C-terminus as previously described (Wells et al., 2019). Constructs for NanoBRET measurements of TTBK1 (NLuc-TTBK1 and TTBK1-NLuc) and TTBK2 (NLuc-TTBK2 and TTBK2-NLuc) were kindly provided by Promega. NLuc orientations used in the respective assays are indicated. Cells were plated in 96-well tissue culture treated plates (Corning 3917) at a cell density of 2 × 10^5^ cells/mL, with a total volume of 100 μL per well in DMEM. After 16 h, the media was aspirated from the plate and replaced with room temperature Opti-MEM without phenol red (Gibco, 100 μL in no tracer wells; 95 μL in tracer only wells; 90 μL in no tracer wells with digitonin; 95 μL in only tracer wells with digitonin). Plates were incubated for 25 min if digitonin was added or for 2 h for the intact cell NanoBRET assay. NanoBRET plates were read after addition of 50 μL of a stock solution (3X) containing NanoBRET NanoGlo substrate (Promega N2161), extracellular NanoLuc inhibitor (Promega N2161) and Opti-MEM. For a 96-well plate the 3X stock solution was prepared with 30 μL of NanoBRET NanoGlo substrate, 10 μL of extracellular NanoLuc inhibitor, and 4,960 μL of Opti-MEM without phenol red. Raw milliBRET units (mBU) were read on a GloMax Discover system with a donor emission wavelength of 450 nm and an acceptor emission wavelength of 600 nm. mBU were calculated by dividing the acceptor emission values (600 nm) by the donor emission values (450 nm). For NanoBRET studies with TTBK1/2, background corrected mBU were calculated by subtracting the no tracer wells from the tracer containing wells and multiplying by 1000.

#### Tracer Permeation Studies

HEK293 cells were transfected with constructs of TTBK1 or TTBK2 tagged with NLuc on the N- or C-terminus. Tracer **29** (20X) was prepared from a 400 μM stock solution in DMSO with tracer dilution buffer (Promega N291B) and 20% DMSO, with a final assay plate concentration of 1% DMSO. 5 μL of **29** (20X) was added to each well, with exception of the no tracer control wells, with final plate concentrations of 0.5, 1, and 2 μM. 10 μL of digitonin (Fisher, 10X) was added to specific wells to permeabilize the cells. The 10X digitonin stock was prepared by diluting a 40X DMSO solution of digitonin with room temperature Opti-MEM without phenol red, for a final assay plate concentration of 1% DMSO. Plates containing digitonin were incubated for 25 min and non-digitonin containing plates were incubated for 2 h. Two biological replicates each with two technical replicates were plotted in a bar chart with the standard deviation represented as error bars (Figure S4).

#### Tracer Competition Analyses

HEK293 cells were transfected with NLuc-TTBK1 or NLuc-TTBK2. Compounds **3, 9**, and **10** were used to compete away the NanoBRET signal produced with tracer **29**. These compounds displace the tracer due to occupancy of the same binding site. Tracer **29** (20X) stock solutions were prepared in tracer dilution buffer (Promega N291B) for final assay plate concentrations of 0.125, 0.25, 0.5, 1, and 2 μM, with a final assay plate concentration of 1% DMSO. 10X stock solutions of compounds **3, 9**, and **10** were prepared from 10 mM DMSO stock solutions with room temperature Opti-MEM without phenol red, for final assay plate concentrations of 10 μM and 30 μM. 5 μL of **29** (20X) was added to each well, with exception of the no tracer control wells, and 10 μL of 10X solutions of compounds **3, 9**, and **10** were added to wells for tracer competition. One biological replicate was plotted in GraphPad Prism with [inhibitor] vs. response (three parameters) and included in Figure 3B.

#### Tracer Titration Experiments

HEK293 cells were transfected with NLuc-TTBK1 or NLuc-TTBK2. Compound **3** was tested in 12-point dose–response format with a top concentration of 30 μM. Tracer **29** (20X) stock solutions were prepared in tracer dilution buffer (Promega N291B) and 20% DMSO, for final assay plate concentrations of 0.125, 0.25, 0.5, 1, and 2 μM. The final assay plate concentrations included 1% DMSO. 5 μL of **29** (20X) was added to each well, with exception of the no tracer control wells, and 10 μL of 10X 3-fold diluted solutions of compound **3** was added to wells. One biological replicate was plotted in GraphPad Prism with log[inhibitor] vs. response (three parameters) and included in Figure 3C.

#### NanoBRET Assays

NanoBRET assays were carried out in dose–response format as described previously (Wells *et al*., 2019). NLuc orientations used in the respective assays are as follows: NLuc-TTBK1, NLuc-TTBK2, and PIP4K2C-NLuc. Based on the tracer titration results, assays were carried as described by the manufacturer using 2.0 μM of tracer **29** for TTBK1, 2.0 μM of tracer **29** for TTBK2, and 0.063 μM of tracer K8 for PIP4K2C. 20X stock solutions of the respective tracers were prepared in tracer dilution buffer (Promega N291B) and 20% DMSO for final assay plate concentration of 2 μM (TTBK1/2) or 0.063 μM (PIP4K2C), with a final assay plate concentration of 1% DMSO. For TTBK1 and TTBK2, compounds were tested in 12-point dose– response format with a top concentration of 30 μM (data in Table 2). For PIP4K2C, compound **9** was tested in 12-point dose–response format with a top concentration of 10 μM (data in Table S2). One biological replicate was plotted in GraphPad Prism with log[inhibitor] vs. response (three parameters). Plots of TTBK1/2 data are included in Figures 3D and S4.

#### SH-SY5Y Western Blot Analyses and Lysosomal Imaging

SH-SY5Y cells were exposed to **9** or **10** for the time period indicated, with or without co-treatment with ethacrynic acid (EA), in Figure 3/S2 before harvesting lysates. 0.05% DMSO was used as a control. Lysates were collected in RIPA, sonicated, and quantified using BCA assay. 20 μg of protein was run on each lane. Blots were probed with 1:5000 phospho-TDP-43 (Proteintech #22309-1-AP) or 1:10,000 TDP-43 (Proteintech #10782-2-AP). 1:5000 Actin (Sigma #A2228) was used as a loading control on each blot. TDP-43 protein was normalized to actin loading control only. Phospho-TDP-43 was normalized to actin, and then normalized to TDP-43 to account for differences in (total) TDP-43 expression between doses.

For imaging of lysosomes (Figure S2), SH-SY5Y cells were plated in 35mm glass bottom dishes and treated for 24 h with either 0.05% DMSO (control) or 5 μM of **9** or **10**. After 24 h, 50 nM LysoTracker Red DND-99 (ThermoFisher) was added to each dish, and they were incubated for 30 minutes at 37°C. Live cells were imaged at 40X magnification using a microscope equipped with a camera. ImageJ software was utilized for image generation.

#### Kinetic Solubility

Analysis of kinetic solubility was carried out by Analiza, Inc using 10 mM DMSO stocks of compounds in phosphate buffered saline solution (PBS) at pH 7.4 as described previously (Drewry *et al*., 2022a). Calculated solubility values in Table 2 have been corrected for background nitrogen present in the DMSO and media.

#### Crystallography and Biophysical Methods

##### Crystallization and Structure Determination

Recombinant TTBK1 (aa 13-320) co-expressed with lambda phosphatase in *E. coli* Rosetta as a His-Sumo-tagged protein was initially purified by Ni2+-affinity chromatography, and the tag was cleaved by SENP1 protease treatment. The cleaved proteins were further purified by size exclusion chromatography and stored in buffer composed of 25 mM HEPES pH 7.5, 250 mM NaCl, 0.5 mM TCEP and 10% glycerol. The kinase at 10 mg/ml was mixed with inhibitors at 1 mM, and the complexes were crystallized using sitting drop vapor diffusion method at 20 °C and the condition containing 14-23% PEG 3350, 0.2 M sodium acetate pH 7.0 and 0.1 M tris pH 7.5–9.0. Crystals were cryo-protected with mother liquor supplemented with 22% ethylene glycol. Diffraction data collected at Swiss Light Source were processed and scaled using XDS (Kabsch, 2010) and Aimless (Evans and Murshudov, 2013), respectively. The structures were solved by molecular replacement using Phaser (McCoy et al., 2007) and the coordinate of TTBK1 (PDB code: 7Q8W (Nozal *et al*., 2022)). Model rebuilding was performed using COOT (Emsley et al., 2010) and refinement via Refmac5 (Murshudov et al., 2011). Solved co-crystal structures are included in Figure 2 and the data collection and refinement statistics are summarized in Table S3.

##### Thermal Shift Assays

Either the TTBK1 or TTBK2 kinase domain at 2 μM in 10 mM HEPES, pH 7.5 and 500 mM NaCl was mixed with the inhibitors at 10 μM in the presence of SyPRO orange dye (Invitrogen). A Real-Time PCR Mx3005p machine (Stratagene) was used to measure the fluorescence spectrum. The Tm shift assays and data evaluation for melting temperatures were performed according to the previously described protocol (Fedorov et al., 2007).

#### Phenotypic Studies

##### Cilia Imaging in iPSCs

Human iPSCs (1 × 10^4^ cells/well) were plated in Matrigel-coated black 96 well plates in culture medium supplemented with 10 μM Y27632 and incubated for two hours at 37°C and 5% CO_2_. Then, Y27632 was removed, and starvation was induced by not changing the culture medium for the next 72 hours. For TTBK inhibitor treatments of human iPSCs, **9** or **10** was introduced at concentrations of 0.1 μM, 0.5 μM or 1 μM when the medium change was performed to remove Y27632 (for 72 h continuous treatment) or after 48 h of starvation (for 24 h continuous treatment). After the starvation and co-treatment period, cells were fixed for 5 min with 4% formaldehyde at room temperature, followed by 5 min with 100% methanol at -20°C, and then permeabilized for 15 min with 0.3% Triton X and blocked with 5% BSA at room temperature. Primary antibodies, anti-ARL13B (Abcam ab153725) and anti-gamma tubulin (Sigma T6557), were used at 1:500 dilution for 2 hours at room temperature. Secondary antibodies (anti-rabbit AlexaFluor 488 and anti-mouse AlexaFluor 647, both from LifeTechnologies) were used at 1:1000 dilution for 30 min at room temperature. Nuclei were counterstained with DAPI (Sigma) for 5 min. Images were taken using the built-in camera of the Life Technologies EVOS FL microscope. Both GFP and DAPI EVOS Light Cubes were used for cilia and nuclei detection. Image quantification was done using ImageJ Fiji by modifying the threshold to select the desired stained phenotype and then minimizing noise and analyzing particles. Six to ten images of random fields were quantified for each sample and results were expressed as the percentage of cells expressing primary cilia.

### QUANTIFICATION AND STATISTICAL ANALYSIS

#### Statistical analysis

The statistical methods used have been indicated under the figures in which error bars or statistics are included. Replicate numbers indicate the number of biological replicates analyzed to generate the summary figures. GraphPad Prism 8.2.0 software was used for analyses unless otherwise indicated in figure legends or experimental protocols.

## REFERENCES

Bao, C., Bajrami, B., Marcotte, D.J., Chodaparambil, J.V., Kerns, H.M., Henderson, J., Wei, R., Gao, B., and Dillon, G.M. (2021). Mechanisms of regulation and diverse activities of tau-tubulin kinase (TTBK) isoforms. Cell Mol. Neurobiol. 41, 669–685.

Battaglia, R.A., Beltran, A.S., Delic, S., Dumitru, R., Robinson, J.A., Kabiraj, P., Herring, L.E., Madden, V.J., Ravinder, N., Willems, E., et al. (2019). Site-specific phosphorylation and caspase cleavage of GFAP are new markers of Alexander disease severity. Elife 8, e47789.

Beltran, A.A., Molina, S.G., Marquez, A., Munoz, L.J., Olivares, J.F., and Beltran, A.S. (2021). Generation of an induced pluripotent stem cell line (UNCCi002-A) from a healthy donor using a non-integration system to study Cerebral Cavernous Malformation (CCM). Stem Cell Res. 54, 102421.

Bosc, N., Meyer, C., and Bonnet, P. (2017). The use of novel selectivity metrics in kinase research. BMC Bioinform. 18, 17.

Bowie, E., and Goetz, S.C. (2020). TTBK2 and primary cilia are essential for the connectivity and survival of cerebellar Purkinje neurons. Elife 9, e51166.

Bowie, E., Norris, R., Anderson, K.V., and Goetz, S.C. (2018). Spinocerebellar ataxia type 11-associated alleles of Ttbk2 dominantly interfere with ciliogenesis and cilium stability. PLoS Genet. 14, e1007844.

Cajánek, L., and Nigg, E.A. (2014). Cep164 triggers ciliogenesis by recruiting Tau tubulin kinase 2 to the mother centriole. Proc. Natl. Acad. Sci. U.S.A. 111, E2841–2850.

Davis, M.I., Hunt, J.P., Herrgard, S., Ciceri, P., Wodicka, L.M., Pallares, G., Hocker, M., Treiber, D.K., and Zarrinkar, P.P. (2011). Comprehensive analysis of kinase inhibitor selectivity. Nat. Biotechnol. 29, 1046–1051.

Dillon, G.M., Henderson, J.L., Bao, C., Joyce, J.A., Calhoun, M., Amaral, B., King, K.W., Bajrami, B., and Rabah, D. (2020). Acute inhibition of the CNS-specific kinase TTBK1 significantly lowers tau phosphorylation at several disease relevant sites. PLoS One 15, e0228771.

Drewry, D. H., Annor-Gyamfi, J. K., Wells, C. I., Pickett, J. E., Dederer, V., Preuss, F., Mathea, S., and Axtman, A. D. (2022a). Identification of pyrimidine-based lead compounds for understudied kinases implicated in driving neurodegeneration. J. Med. Chem. 65, 1313–1328.

Drewry, D.H., Potjewyd, F.M., Smith, J.L., Dickmander, R.J., Bayati, A., Howell, S., Taft-Benz, S.A., Hossain, M.A., Heise, M.T., McPherson, P.S., et al. (2022b). Identification and utilization of a chemical probe to interrogate the roles of PIKfyve in the lifecycle of β-coronaviruses. ChemRxiv. 10.26434/chemrxiv-2022-bj274.

Drewry, D.H., Wells, C.I., Andrews, D.M., Angell, R., Al-Ali, H., Axtman, A.D., Capuzzi, S.J., Elkins, J.M., Ettmayer, P., Frederiksen, M., et al. (2017). Progress towards a public chemogenomic set for protein kinases and a call for contributions. PLoS One 12, e0181585.

Emsley, P., Lohkamp, B., Scott, W.G., and Cowtan, K. (2010). Features and development of Coot. Acta Crystallogr. D Biol. Crystallogr. 66, 486–501.

Evans, P.R., and Murshudov, G.N. (2013). How good are my data and what is the resolution? Acta Crystallogr. D Biol. Crystallogr. 69, 1204–1214.

Fedorov, O., Marsden, B., Pogacic, V., Rellos, P., Müller, S., Bullock, A.N., Schwaller, J., Sundström, M., and Knapp, S. (2007). A systematic interaction map of validated kinase inhibitors with Ser/Thr kinases. Proc. Natl. Acad. Sci. U.S.A. 104, 20523–20528.

Gayle, S., Landrette, S., Beeharry, N., Conrad, C., Hernandez, M., Beckett, P., Ferguson, S.M., Mandelkern, T., Zheng, M., Xu, T., et al. (2017). Identification of apilimod as a first-in-class PIKfyve kinase inhibitor for treatment of B-cell non-Hodgkin lymphoma. Blood 129, 1768–1778.

Goetz, S.C., Liem, K.F., Jr., and Anderson, K.V. (2012). The spinocerebellar ataxia-associated gene Tau tubulin kinase 2 controls the initiation of ciliogenesis. Cell 151, 847–858.

Halkina, T., Henderson, J.L., Lin, E.Y., Himmelbauer, M.K., Jones, J.H., Nevalainen, M., Feng, J., King, K., Rooney, M., Johnson, J.L., et al. (2021). Discovery of potent and brain-penetrant tau tubulin kinase 1 (TTBK1) inhibitors that lower tau phosphorylation in vivo. J. Med. Chem. 64, 6358–6380.

Houlden, H., Johnson, J., Gardner-Thorpe, C., Lashley, T., Hernandez, D., Worth, P., Singleton, A.B., Hilton, D.A., Holton, J., Revesz, T., et al. (2007). Mutations in TTBK2, encoding a kinase implicated in tau phosphorylation, segregate with spinocerebellar ataxia type 11. Nat. Genet. 39, 1434–1436.

Iguchi, Y., Katsuno, M., Takagi, S., Ishigaki, S., Niwa, J., Hasegawa, M., Tanaka, F., and Sobue, G. (2012). Oxidative stress induced by glutathione depletion reproduces pathological modifications of TDP-43 linked to TDP-43 proteinopathies. Neurobiol. Dis. 45, 862–870.

Ikezu, S., and Ikezu, T. (2014). Tau-tubulin kinase. Front. Mol. Neurosci. 7, 33.

Ikezu, S., Ingraham Dixie, K.L., Koro, L., Watanabe, T., Kaibuchi, K., and Ikezu, T. (2020). Tau-tubulin kinase 1 and amyloid-β peptide induce phosphorylation of collapsin response mediator protein-2 and enhance neurite degeneration in Alzheimer disease mouse models. Acta Neuropathol. Commun. 8, 12.

Kabsch, W. (2010). XDS. Acta Crystallogr. D Biol. Crystallogr. 66, 125–132.

Li, K., McGee, L.R., Fisher, B., Sudom, A., Liu, J., Rubenstein, S.M., Anwer, M.K., Cushing, T.D., Shin, Y., Ayres, M., et al. (2013). Inhibiting NF-κB-inducing kinase (NIK): Discovery, structure-based design, synthesis, structure–activity relationship, and co-crystal structures. Bioorg. Med. Chem. Lett. 23, 1238–1244.

Liachko, N.F., McMillan, P.J., Strovas, T.J., Loomis, E., Greenup, L., Murrell, J.R., Ghetti, B., Raskind, M.A., Montine, T.J., Bird, T.D., et al. (2014). The tau tubulin kinases TTBK1/2 promote accumulation of pathological TDP-43. PLoS Genet. 10, e1004803.

Liao, J.-C., Yang, T.T., Weng, R.R., Kuo, C.-T., and Chang, C.-W. (2015). TTBK2: A tau protein kinase beyond tau phosphorylation. Biomed. Res. Int. 2015, 575170.

Lo, C.H., Lin, I.H., Yang, T.T., Huang, Y.C., Tanos, B.E., Chou, P.C., Chang, C.W., Tsay, Y.G., Liao, J.C., and Wang, W.J. (2019). Phosphorylation of CEP83 by TTBK2 is necessary for cilia initiation. J. Cell. Biol. 218, 3489–3505.

Lund, H., Cowburn, R.F., Gustafsson, E., Stromberg, K., Svensson, A., Dahllund, L., Malinowsky, D., and Sunnemark, D. (2013). Tau-tubulin kinase 1 expression, phosphorylation and co-localization with phospho-Ser422 tau in the Alzheimer’s disease brain. Brain Pathol. 23, 378–389.

Ma, R., Kutchy, N.A., Chen, L., Meigs, D.D., and Hu, G. (2022). Primary cilia and ciliary signaling pathways in aging and age-related brain disorders. Neurobiol. Dis. 163, 105607.

Marcotte, D.J., Spilker, K.A., Wen, D., Hesson, T., Patterson, T.A., Kumar, P.R., and Chodaparambil, J.V. (2020). The crystal structure of the catalytic domain of tau tubulin kinase 2 in complex with a small-molecule inhibitor. Acta Crystallogr. F 76, 103–108.

May, E.A., Sroka, T.J., and Mick, D.U. (2021). Phosphorylation and ubiquitylation regulate protein trafficking, signaling, and the biogenesis of primary cilia. Front. Cell. Dev. Biol. 9, 664279–664279.

McCoy, A.J., Grosse-Kunstleve, R.W., Adams, P.D., Winn, M.D., Storoni, L.C., and Read, R.J. (2007). Phaser crystallographic software. J. Appl. Crystallogr. 40, 658–674.

McMillan, P., Wheeler, J., Gatlin, R.E., Taylor, L., Strovas, T., Baum, M., Bird, T.D., Latimer, C., Keene, C.D., Kraemer, B.C., and Liachko, N.F. (2020). Adult onset pan-neuronal human tau tubulin kinase 1 expression causes severe cerebellar neurodegeneration in mice. Acta Neuropathol. Commun. 8, 200.

Molina, S.G., Beltran, A.A., and Beltran, A.S. (2020). Generation of an integration-free induced pluripotent stem cell line (UNC001-A) from blood of a healthy individual. Stem Cell Res. 49, 102015.

Murshudov, G.N., Skubák, P., Lebedev, A.A., Pannu, N.S., Steiner, R.A., Nicholls, R.A., Winn, M.D., Long, F., and Vagin, A.A. (2011). REFMAC5 for the refinement of macromolecular crystal structures. Acta Crystallogr. D Biol. Crystallogr. 67, 355–367.

Nozal, V., and Martinez, A. (2019). Tau tubulin kinase 1 (TTBK1), a new player in the fight against neurodegenerative diseases. Eur. J. Med. Chem. 161, 39–47.

Nozal, V., Martínez-González, L., Gomez-Almeria, M., Gonzalo-Consuegra, C., Santana, P., Chaikuad, A., Pérez-Cuevas, E., Knapp, S., Lietha, D., Ramírez, D., et al. (2022). TDP-43 modulation by tau-tubulin kinase 1 inhibitors: A new avenue for future amyotrophic lateral sclerosis therapy. J. Med. Chem. 65, 1585–1607.

Oda, T., Chiba, S., Nagai, T., and Mizuno, K. (2014). Binding to Cep164, but not EB1, is essential for centriolar localization of TTBK2 and its function in ciliogenesis. Genes Cells 19, 927–940.

Robers, M.B., Dart, M.L., Woodroofe, C.C., Zimprich, C.A., Kirkland, T.A., Machleidt, T., Kupcho, K.R., Levin, S., Hartnett, J.R., Zimmerman, K., et al. (2015). Target engagement and drug residence time can be observed in living cells with BRET. Nat. Commun. 6, 10091.

Rodgers, G., Austin, C., Anderson, J., Pawlyk, A., Colvis, C., Margolis, R., and Baker, J. (2018). Glimmers in illuminating the druggable genome. Nat. Rev. Drug Discov. 17, 301–302.

Sánchez, I., and Dynlacht, B.D. (2016). Cilium assembly and disassembly. Nat. Cell Biol. 18, 711–717.

Sato, S., Cerny, R.L., Buescher, J.L., and Ikezu, T. (2006). Tau-tubulin kinase 1 (TTBK1), a neuron-specific tau kinase candidate, is involved in tau phosphorylation and aggregation. J. Neurochem. 98, 1573–1584.

Sato, S., Xu, J., Okuyama, S., Martinez, L.B., Walsh, S.M., Jacobsen, M.T., Swan, R.J., Schlautman, J.D., Ciborowski, P., and Ikezu, T. (2008). Spatial learning impairment, enhanced CDK5/p35 activity, and downregulation of NMDA receptor expression in transgenic mice expressing tau-tubulin kinase 1. J. Neurosci. 28, 14511–14521.

Vasta, J.D., Corona, C.R., Wilkinson, J., Zimprich, C.A., Hartnett, J.R., Ingold, M.R., Zimmerman, K., Machleidt, T., Kirkland, T.A., Huwiler, K.G., et al. (2018). Quantitative, wide-spectrum kinase profiling in live cells for assessing the effect of cellular ATP on target engagement. Cell Chem. Biol. 25, 206–214.

Wells, C., Couñago, R.M., Limas, J.C., Almeida, T.L., Cook, J.G., Drewry, D.H., Elkins, J.M., Gileadi, O., Kapadia, N.R., Lorente-Macias, A., et al. (2019). SGC-AAK1-1: A chemical probe targeting AAK1 and BMP2K. ACS Med. Chem. Lett. 11, 340–345.

Xue, Y., Wan, P.T., Hillertz, P., Schweikart, F., Zhao, Y., Wissler, L., and Dekker, N. (2013). X-ray structural analysis of tau-tubulin kinase 1 and its interactions with small molecular inhibitors. ChemMedChem 8, 1846–1854.

Yu, N.N., Yu, J.T., Xiao, J.T., Zhang, H.W., Lu, R.C., Jiang, H., Xing, Z.H., and Tan, L. (2011). Tau-tubulin kinase-1 gene variants are associated with Alzheimer’s disease in Han Chinese. Neurosci. Lett. 491, 83–86.

