## Supplementary material for "Modulation of Tau Tubulin Kinases (TTBK1 and TTBK2) Impacts Ciliogenesis": Combined Supplemental Data

### SELECTIVITY SCREENING, CRYSTALLOGRAPHIC DETAILS, SYNTHETIC ROUTES, CELL-BASED ASSAY RESULTS, and GENETIC EDITING DETAILS

#### Table S1. Enzymatic Profiling of Indolyl Pyrimidinamine Library in Targeted Kinase Panel, Related to Compounds in Tables 1 and 2

| Compound | % control at 1 μM[a] | | | | | | | | | | | | | | | | |
|---|---|---|---|---|---|---|---|---|---|---|---|---|---|---|---|---|---|
| | HIPK1[b] | MAP4K5 | MARK3[b] | MARK4[b] | PAK3[b] | PAK4 | PAK5[b] | PAK6[b] | TAOK1[b] | TAOK2[b] | TAOK3 | TSSK1[b] | TSSK2[b] | TSSK3[b] | TSSK4[b] | TTBK1[b] | TTBK2[b] |
| **1** | 76 | 3 | 56 | 53 | 86 | 58 | 48 | 51 | 5 | 28 | 11 | 48 | 88 | 11 | 55 | 78 | 43 |
| **AMG28** | 67 | 1 | 12 | 18 | 86 | 8 | 3 | 6 | 2 | 26 | 9 | 10 | 57 | 1 | 70 | 50 | 20 |
| **2** | 96 | 9 | 76 | 86 | 100 | 67 | 87 | 77 | 65 | 83 | 78 | 82 | 76 | 120 | 77 | 92 | 69 |
| **3** | 87 | 1 | 46 | 43 | 104 | 30 | 31 | 40 | 7 | 12 | 20 | 30 | 78 | 5 | 78 | 43 | 24 |
| **4** | 77 | 2 | 52 | 48 | 81 | 21 | 27 | 18 | 11 | 33 | 20 | 46 | 82 | 24 | 75 | 78 | 64 |
| **5** | 69 | -1 | 27 | 37 | 89 | 10 | 8 | 13 | 2 | 8 | 9 | 8 | 55 | 13 | 86 | 68 | 72 |
| **6** | 30 | 1 | 31 | 37 | 89 | 22 | 26 | 36 | 1 | 9 | 11 | 57 | 91 | 36 | 103 | 91 | 76 |
| **7** | 56 | 3 | 84 | 84 | 110 | 31 | 39 | 40 | 24 | 21 | 28 | 98 | 93 | 34 | 103 | 86 | 104 |
| **8** | 84 | 6 | 83 | 73 | 117 | 66 | 77 | 82 | 22 | 55 | 44 | 73 | 77 | 59 | 94 | 85 | 103 |
| **9** | 124 | 2 | 77 | 60 | 103 | 36 | 60 | 40 | 36 | 31 | 43 | 38 | 76 | 5 | 102 | 41 | 17 |
| **10** | 89 | 1 | 48 | 39 | 117 | 12 | 17 | 14 | 9 | 14 | 18 | 16 | 58 | -3 | 88 | 37 | 18 |
| **11** | 96 | 9 | 104 | 103 | 100 | 91 | 88 | 92 | 73 | 44 | 89 | 89 | 86 | 75 | 101 | 117 | 110 |

[a] Compounds tested at a single concentration (1 μM) in duplicate. [b] IDG kinase

#### Table S2. Enzymatic and NanoBRET Follow-up of Kinases Potently Inhibited by 9, Related to Selectivity Data in Tables 1

| Kinase[a] | DiscoverX PoC value | Assay format | $IC_{50}$ (nM) |
|---|---|---|---|
| CHK2 | 22 | Enzymatic | 4060 |
| CLK2 | 27 | Enzymatic | 2750 |
| CRIK | 30 | Enzymatic | >10000 |
| CSNK1D | 21 | Enzymatic | 418 |
| CSNK1E | 13 | Enzymatic | 824 |
| MAP4K5 | 21 | Enzymatic | 6 |
| MKK4 | 3.5 | Enzymatic | >10000 |
| MYLK4[b] | 2.4 | Enzymatic | 89.9 |
| PIKfyve[c] | 0 | Enzymatic | 1.77 |
| PIP4K2C[d] | 15 | NanoBRET | 576 |
| PIP5K1C | 0 | Enzymatic | 2060 |
| PKD3 | 0 | Enzymatic | 1120 |
| RIPK5[b] | 4.1 | Enzymatic | 1310 |
| STK16 | 28 | Enzymatic | 297 |
| TAOK1 | 24 | Enzymatic | 782 |
| TAOK2 | 66 | Enzymatic | 862 |
| TSSK3 | 24 | Enzymatic | 235 |
| TTBK1 | ND[e] | Enzymatic | 384 |
| TTBK2 | ND | Enzymatic | 175 |
| YSK4[b] | 1.1 | Enzymatic | 50.6 |

[a] All kinase assays executed at Eurofins unless noted. [b] Evaluated by RBC. [c] Evaluated by SignalChem. [d] Evaluated at SGC-UNC. [e] Not determined.

**Table S3. Crystallographic Data Collection and Refinement Statistics, Related to Figure 2**

| **Complex** | TTBK1-AMG28 | TTBK1-3 | TTBK1-9 | TTBK1-10 |
|---|---|---|---|---|
| **PDB accession code** | 7ZHN | 7ZHO | 7ZHP | 7ZHQ |
| ***Data Collection*** | | | | |
| Resolution[a] (Å) | 48.40-1.85 (1.91-1.85) | 47.48-2.08 (2.15-2.08) | 48.47-1.80 (1.86-1.80) | 48.33-1.80 (1.86-1.80) |
| Spacegroup | *C* 2 | $P\ 2_12_12_1$ | *C* 2 | *C* 2 |
| Cell dimensions | *a* = 171.8, *b* = 39.6, *c* = 49.7 Å<br>*α, γ* =90.0°, *β* = 103.2° | *a* = 39.4, *b* = 49.4, *c* = 171.1 Å<br>*α, β, γ* =90.0° | *a* = 172.3, *b* = 39.8, *c* = 49.8 Å<br>*α, γ* =90.0°, *β* = 103.2° | *a* = 171.3, *b* = 39.5, *c* = 49.7 Å<br>*α, γ* =90.0°, *β* = 103.2° |
| No. unique reflections[a] | 28,144 (2,758) | 20,721 (1,914) | 30,838 (2,992) | 30,388 (2,961) |
| Completeness[a] (%) | 100.0 (99.9) | 99.0 (96.9) | 100.0 (100.0) | 100.0 (100.0) |
| I/σI[a] | 13.4 (1.9) | 12.5 (2.1) | 15.2 (2.0) | 13.4 (2.0) |
| $R_{merge}$[a] | 0.086 (0.884) | 0.145 (0.998) | 0.071 (0.852) | 0.085 (0.780) |
| CC (1/2) | 0.999 (0.773) | 0.998 (0.740) | 0.999 (0.817) | 0.999 (0.829) |
| Redundancy[a] | 6.7 (6.4) | 11.7 (9.5) | 6.7 (6.8) | 6.7 (6.4) |
| ***Refinement*** | | | | |
| No. atoms in refinement (P/L/O)[b] | 2,418/ 25/ 225 | 2,419/ 22/ 172 | 2,399/ 27/ 238 | 2,415/ 26/ 224 |
| B factor (P/L/O)[b] ($Å^2$) | 32/ 21/ 38 | 37/ 30/ 39 | 33/ 23/ 39 | 27/ 19/ 33 |
| $R_{fact}$ (%) | 19.0 | 19.6 | 17.7 | 17.4 |
| $R_{free}$ (%) | 21.5 | 25.8 | 21.8 | 21.2 |
| rms deviation bond[c] (Å) | 0.014 | 0.012 | 0.013 | 0.014 |
| rms deviation angle[c] (°) | 1.5 | 1.3 | 1.4 | 1.5 |

[a] Values in brackets show the statistics for the highest resolution shells. [b] P/O indicate protein, ligand and others (water and solvent molecules), respectively. [c] rms indicates root-mean-square.

A

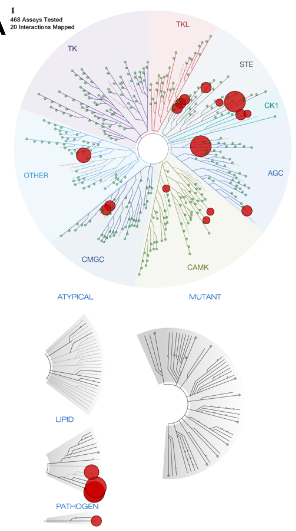


| Kinase | % control at 1000nM |
|---|---|
| YSK4 | 0 |
| PIKFYVE | 0 |
| CIT | 0 |
| PIP5K2C | 0 |
| HASPIN | 0.3 |
| PIP5K1C | 0.5 |
| TAOK1 | 1.2 |
| TAOK3 | 1.5 |
| TAOK2 | 1.5 |
| PKNB(M.tuberculosis) | 1.7 |
| MAP4K2 | 2 |
| CSNK1D | 3 |
| PRKCE | 3.1 |
| CDK11 | 4.7 |
| CDK8 | 4.8 |
| CSNK1E | 6.2 |
| MEK4 | 7.3 |
| RPS6KA4(Kin.Dom.2-C-terminal) | 7.3 |
| CHEK2 | 7.9 |
| MYLK4 | 9.4 |

B

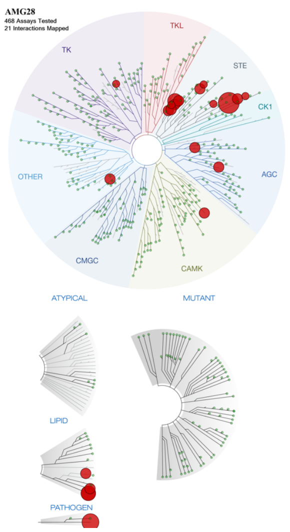


| Kinase | % control at 1000nM |
|---|---|
| YSK4 | 0 |
| PKNB(M.tuberculosis) | 0.1 |
| PAK7 | 0.3 |
| PIKFYVE | 0.4 |
| MAP4K2 | 1.3 |
| CIT | 1.3 |
| TAOK3 | 1.8 |
| RIPK5 | 2.2 |
| MYLK4 | 2.8 |
| PAK6 | 2.9 |
| PIP5K2C | 3.1 |
| PIP5K1C | 3.1 |
| TAOK1 | 3.2 |
| TAOK2 | 3.5 |
| MAP3K2 | 3.7 |
| PRKX | 4.3 |
| PAK4 | 5.7 |
| MAP4K5 | 6.6 |
| JAK2(JH1domain-catalytic) | 6.9 |
| MAP3K3 | 7.2 |
| MEK4 | 8.6 |

C

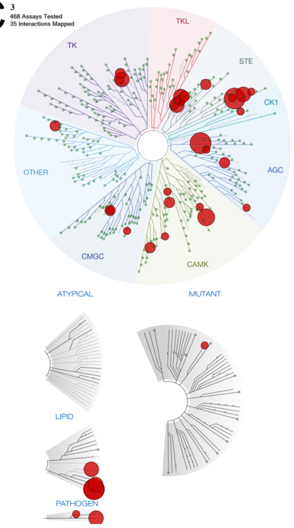


| Kinase | % control at 1000nM |
|---|---|
| YSK4 | 0 |
| PIKFYVE | 0 |
| CIT | 0 |
| RPS6KA4(Kin.Dom.2-C-terminal) | 0.1 |
| PIP5K2C | 0.2 |
| PAK7 | 0.4 |
| TAOK3 | 0.4 |
| PIP5K1C | 0.4 |
| JAK2(JH1domain-catalytic) | 0.5 |
| MAP3K2 | 0.8 |
| PKNB(M.tuberculosis) | 0.9 |
| TAOK2 | 1.3 |
| PRKD3 | 1.3 |
| MAP4K2 | 1.6 |
| SNARK | 1.7 |
| JAK3(JH1domain-catalytic) | 1.8 |
| TAOK1 | 2.1 |
| PRKX | 3.5 |
| AURKB | 3.7 |
| CDC2L1 | 4.3 |
| TYK2(JH1domain-catalytic) | 6.3 |
| BRAF(V600E) | 6.5 |
| CDC2L2 | 6.6 |
| PAK6 | 6.7 |
| PRKCE | 6.8 |
| MAP3K3 | 7.6 |
| PFCDPK1(P.falciparum) | 7.8 |
| TSSK3 | 8.1 |
| PAK4 | 8.3 |
| PRKD2 | 8.3 |
| CSNK1D | 8.4 |
| RPS6KA5(Kin.Dom.2-C-terminal) | 8.9 |
| GRK2 | 9.1 |
| NIK | 9.4 |
| HIPK1 | 9.4 |

D

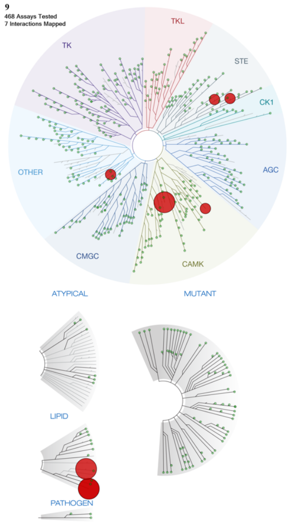


| Kinase | % control at 1000nM |
|---|---|
| PIKFYVE | 0 |
| PIP5K1C | 0 |
| PRKD3 | 0 |
| YSK4 | 1.1 |
| MYLK4 | 2.4 |
| MEK4 | 3.5 |
| RIPK5 | 4.1 |

E

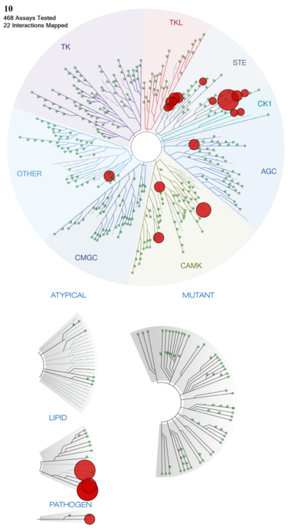


| Kinase | % control at 1000nM |
|---|---|
| PIKFYVE | 0 |
| PIP5K1C | 0 |
| YSK4 | 0 |
| MYLK4 | 0.6 |
| PIP5K2C | 0.6 |
| PAK7 | 1.8 |
| MAP3K2 | 2.2 |
| CIT | 2.4 |
| RIPK5 | 2.4 |
| TSSK3 | 2.4 |
| TAOK1 | 3.1 |
| PAK4 | 4.3 |
| CHEK2 | 4.4 |
| PKNB(M.tuberculosis) | 4.4 |
| MEK4 | 5.5 |
| MAP3K3 | 6.1 |
| PAK6 | 6.7 |
| TAOK3 | 7.8 |
| TAOK2 | 7.9 |
| CSNK1D | 8.2 |
| CSNK1E | 8.9 |
| MAP4K5 | 9.6 |

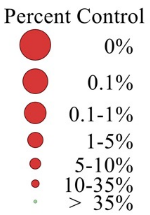


**Figure S1. Kinases that Bind with PoC <10 at 1 μM in *scan*MAX Assay Panel at DiscoverX, Related to Selectivity Data in Table 1**
(A) **1**, (B) AMG28, (C) **3**, (D) **9**, and (E) **10**.

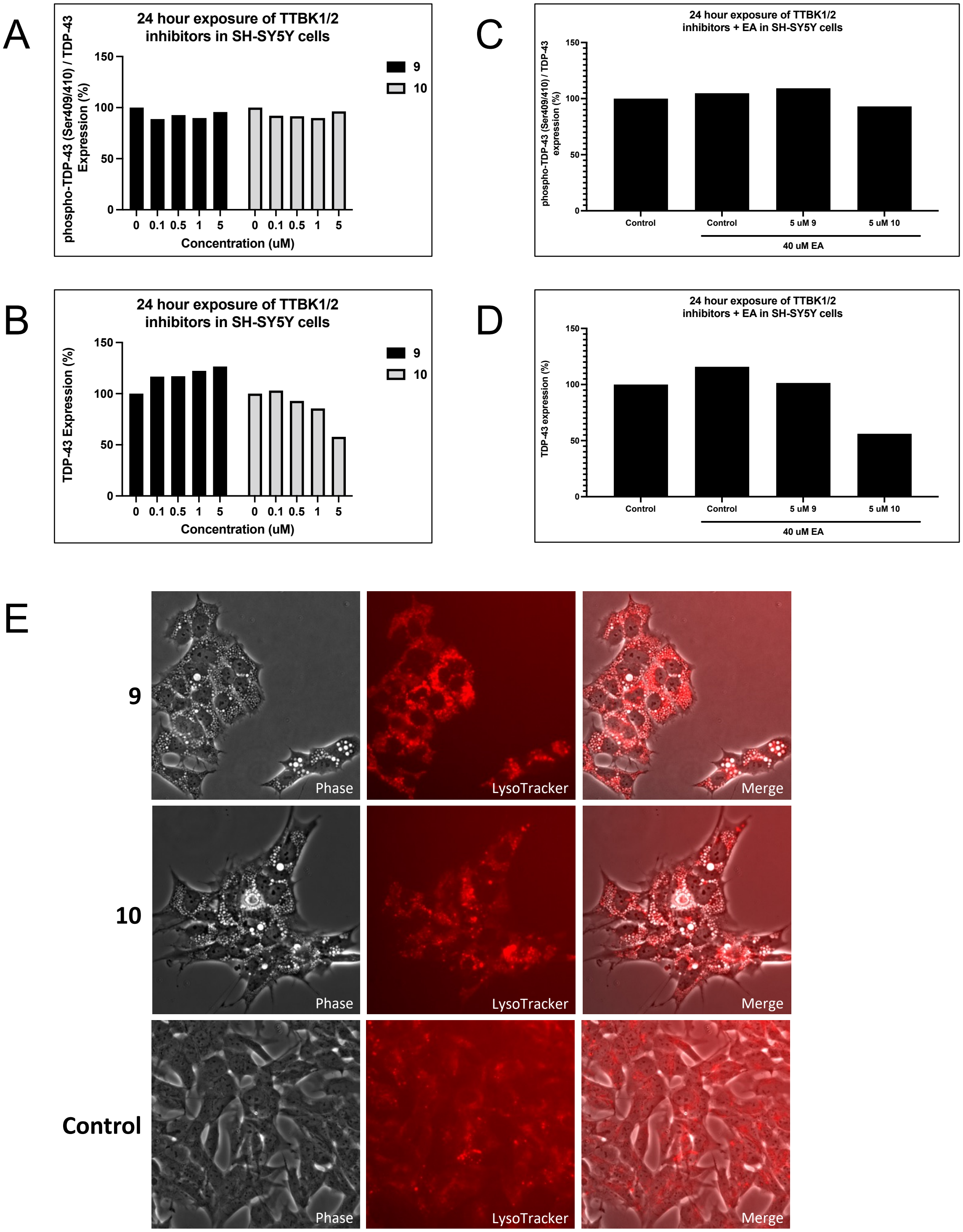


**Figure S2. Western blot Analyses of SH-SY5Y cells after 24 h Treatment with 9 or 10 with and without the Addition of EA, Related to Figure 3**
(A) Quantification of phospho-TDP-43 (Ser409/410, normalized to total TDP-43) after treatment in dose-response, n = 1. (B) Quantification of total TDP-43 after treatment in dose-response, n = 1. (C) Quantification of phospho-TDP-43 (Ser409/410, normalized to total TDP-43) after treatment with 5 μM compound and 40 μM EA, n = 1. (D) Quantification of total TDP-43 after treatment with 5 μM compound and 40 μM EA, n = 1. (E) 50 nM LysoTracker was added after 24 h treatment with 5 uM **9** or **10** and imaged at 40X.

A

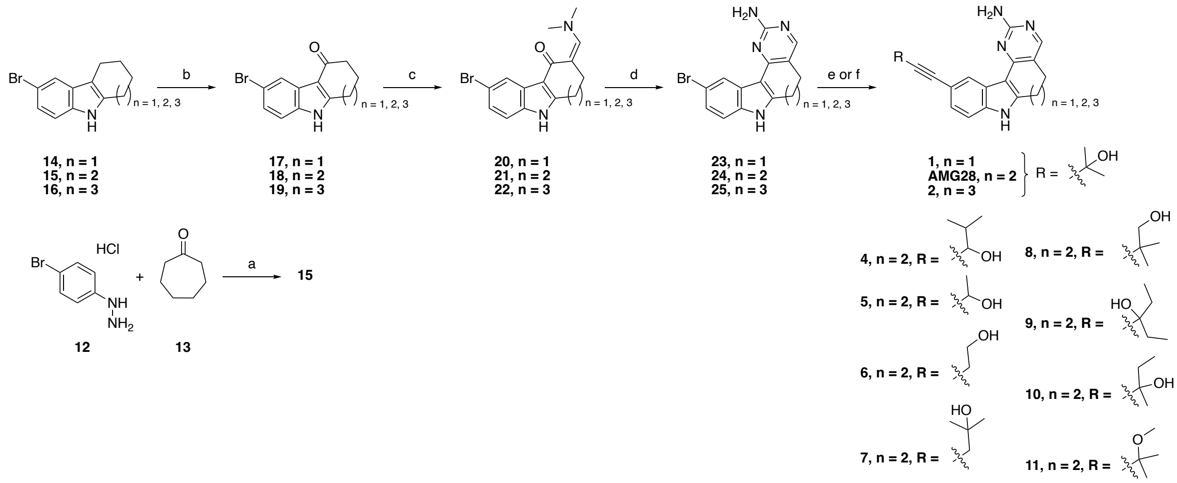


Reagents and conditions: (a) AcOH, 50 °C to 120 °C, 3 h, 70%; (b) DDQ, THF/water, 0 °C, 45 min, 53–87%; (c) Bredereck's reagent, toluene or neat, 110 °C, 3–12 h; (d) NaOMe, guanidine hydrochloride, iPrOH, 100 °C, 12 h, 12–38% over 2 steps; (e) For **1**, **2**, **4**, and AMG28: Alkyne R, TEA, $Pd(PPh_3)_4$, CuI, DMF or DMA, 100 °C, 12 h, 1.6–19%; (f) For **5**–**11**: Alkyne R, DIPA, $PdCl_2(PPh_3)_2$, CuI, propanol, 100 °C, 12 h, 2.4–15%.

B

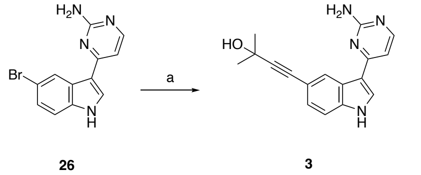


Reagents and conditions: (a) 2-methylbut-3-yn-2-ol, TEA, $Pd(PPh_3)_4$, CuI, DMA, 100 °C, 12 h, 4.4%.

C

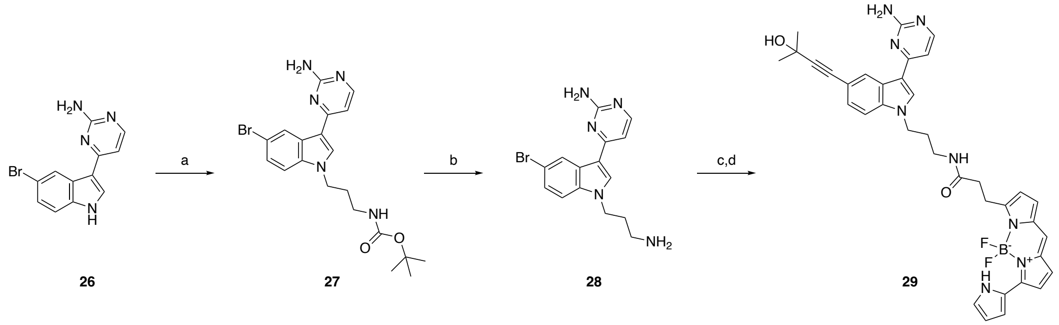


Reagents and conditions: (a) Tos(CH2)$_3$NHBoc, $Cs_2CO_3$, DMF, 50 °C, 16 h, 35%; (b) TFA, DCM, 2 h, 95%; (c) 2-methylbut-3-yn-2-ol, $Pd(PPh_3)_4$, diisopropylamine, CuI, 80 °C, 16 h; (d) NanoBRET590 SE, DIPEA, DMF, 30 min, 0.9% over two steps.

**Figure S3. Schemes Corresponding to Synthesis of all Analogs and Tracer 29, Related to Tables 1 and 2 and Figure 3**

(A) Route and reagents used to prepare analogs **1**, **2**, **4**–**11**, and AMG28. (B) Route and reagents used to prepare analog **3**.

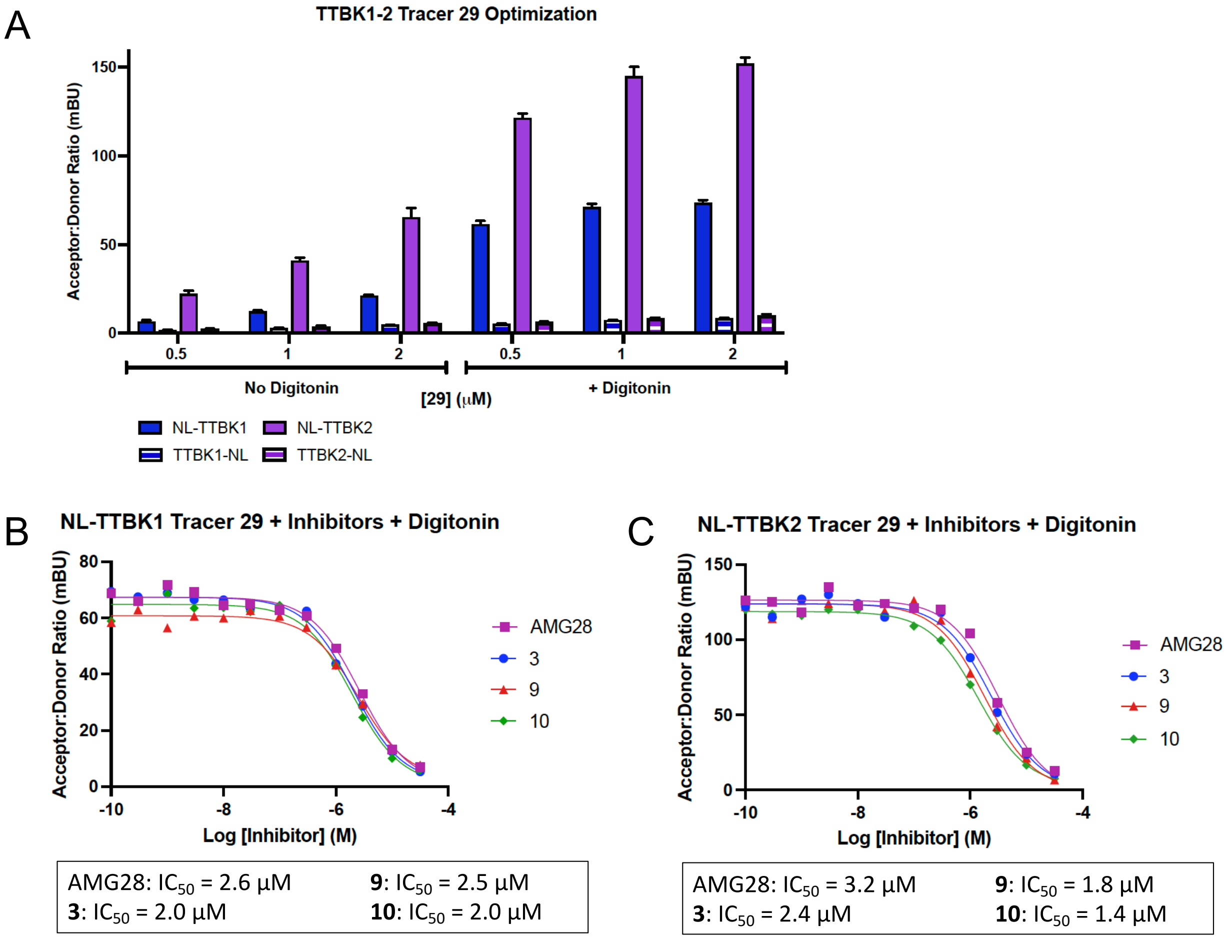


**Figure S4. NanoBRET Data Exploring Cell Permeability of Tracer 29, Related to Figure 3 and Table 2**

(A) Permeation studies for tracer **29** in the presence and absence of digitonin and identification of the optimal NLuc-tagged construct for TTBK1 and TTBK2, n = 2. Error bars represent standard deviation.

(B) Representative NL-TTBK1 NanoBRET assay curves and corresponding $IC_{50}$ values for active compounds AMG28, **3**, **9**, and **10** in permeabilized HEK293 cells, n = 1. (C) Representative NL-TTBK2 NanoBRET assay curves and corresponding $IC_{50}$ values for active compounds AMG28, **3**, **9**, and **10** in permeabilized HEK293 cells, n = 1.

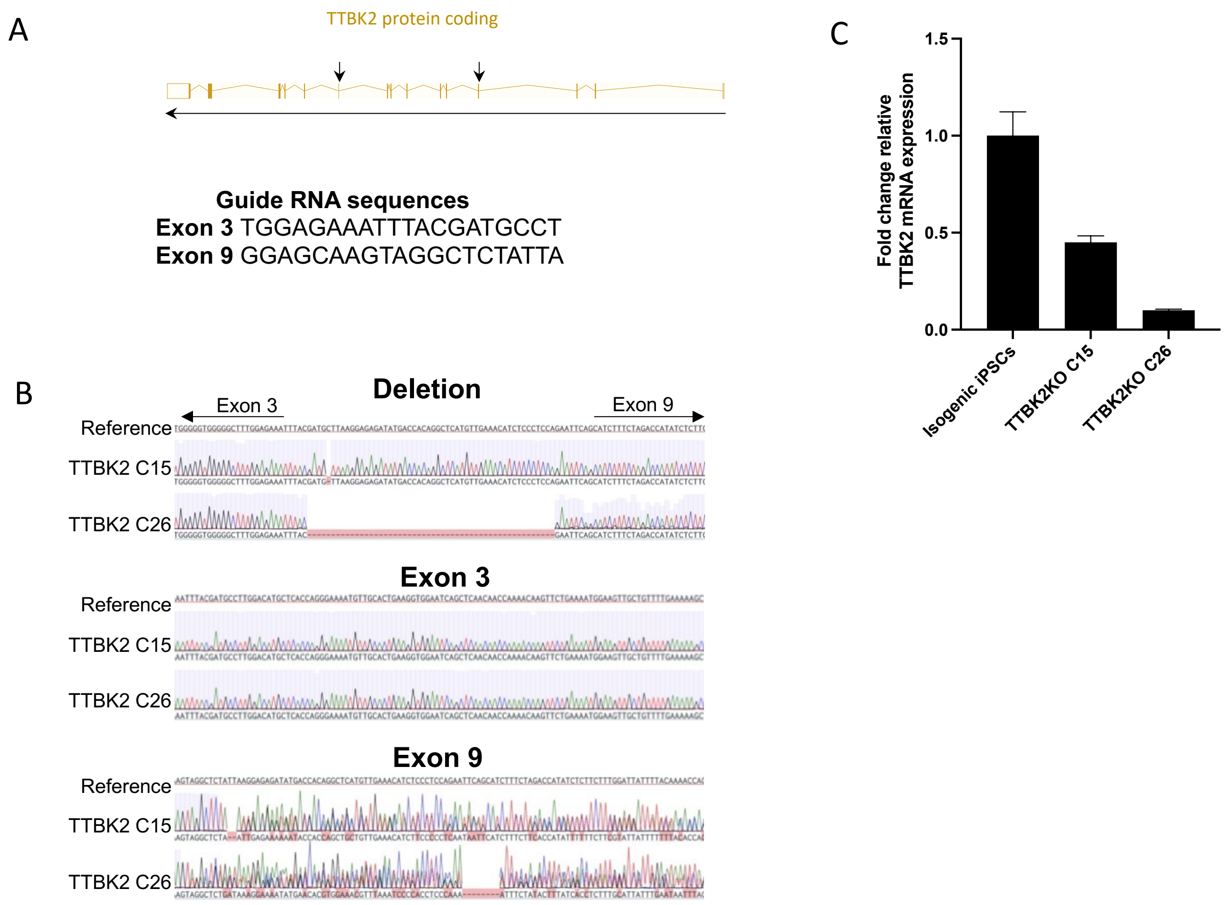


**Figure S5. Generation TTBK2 Knockout iPSCs using CRISPR/Cas9, Related to Figure 5**
(A) Illustration of the genomic context displaying the exons of the TTBK2 gene. Black arrows indicate the exons targeted by CRISPR guide RNAs. (B) Sanger sequencing illustrating the genomic deletion of TTBK2 in two clones (C15 and C26). Sanger sequencing for exon 3, exon 9, and expected deletion in C15 and C26 compared to the isogenic cells (reference). (C) Graph of the TTBK2 mRNA expression levels in C15 and C26 clones relative to the isogenic iPSCs. Data are represented as mean ± SEM.

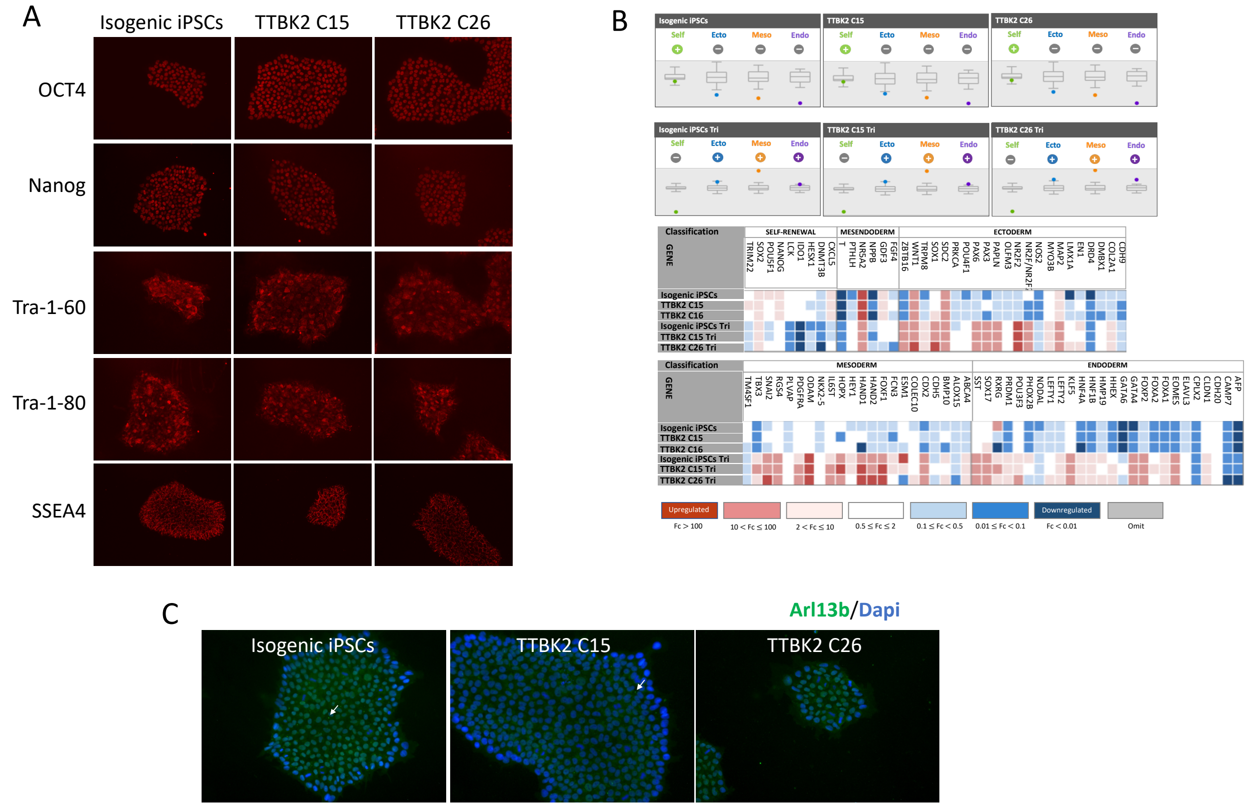


**Figure S6. Characterization of TTBK2 Edited Clones, Related to Figure 5**
(A) Immunofluorescent staining of pluripotency factors OCT4, NANOG, Tra-1-60, Tra-1-80, and SSEA4 in edited iPSCs. (B) Edited TTBK2 cells remain pluripotent (top panel) and can differentiate into the three germ layers using the qRT-PCR based TaqMan hPSC Scorecard Panel assay. (C) Primary cilia expression in TTBK2 edited clones without starvation.

**Data S1: $^1$H Spectra for Final Compounds, Related to Tables 1 and 2 and Figure 3**

$^1$H NMR (400 MHz, MeOD-$d_4$) 4-(2-amino-5,6,7,8-tetrahydropyrimido[4',5':3,4]cyclohepta[1,2-b]indol-11-yl)-2-methylbut-3-yn-2-ol (AMG28):

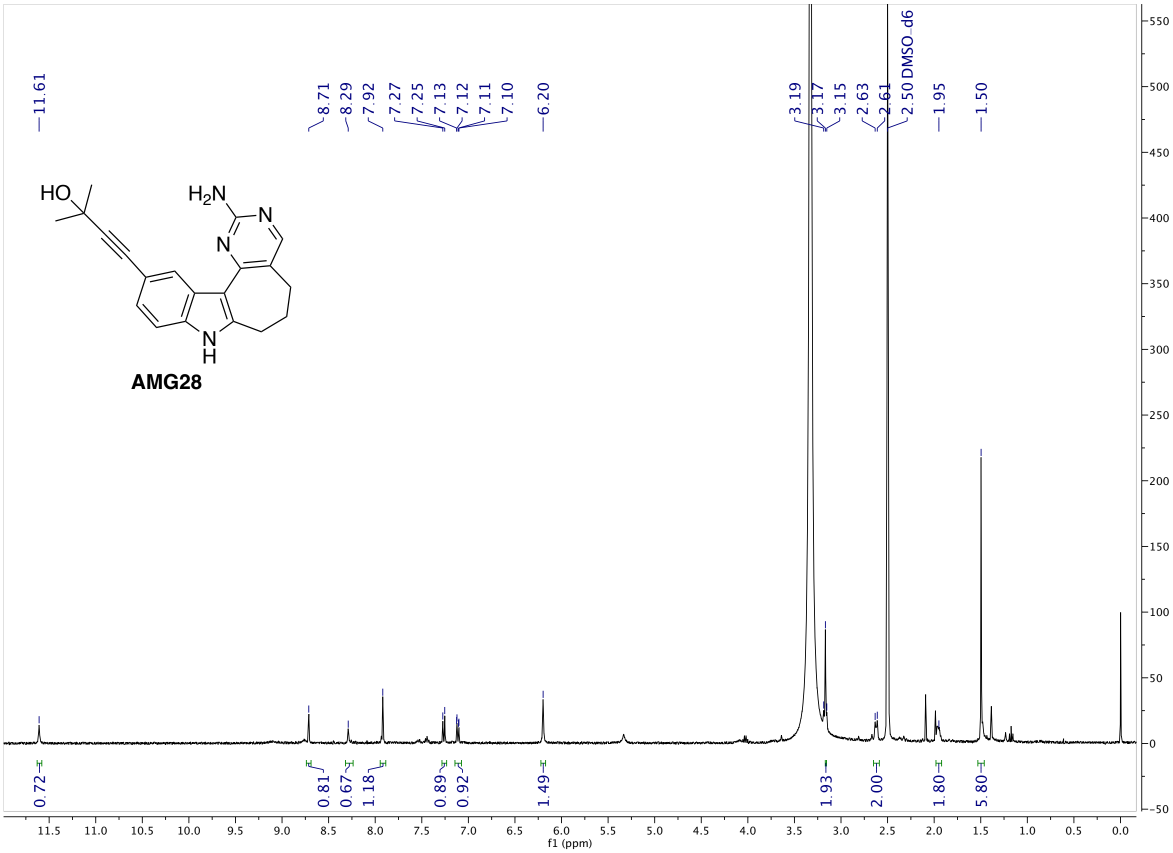

$^1$H NMR (400 MHz, MeOD-$d_4$) 4-(3-(2-aminopyrimidin-4-yl)-1H-indol-5-yl)-2-methylbut-3-yn-2-ol (**3**):

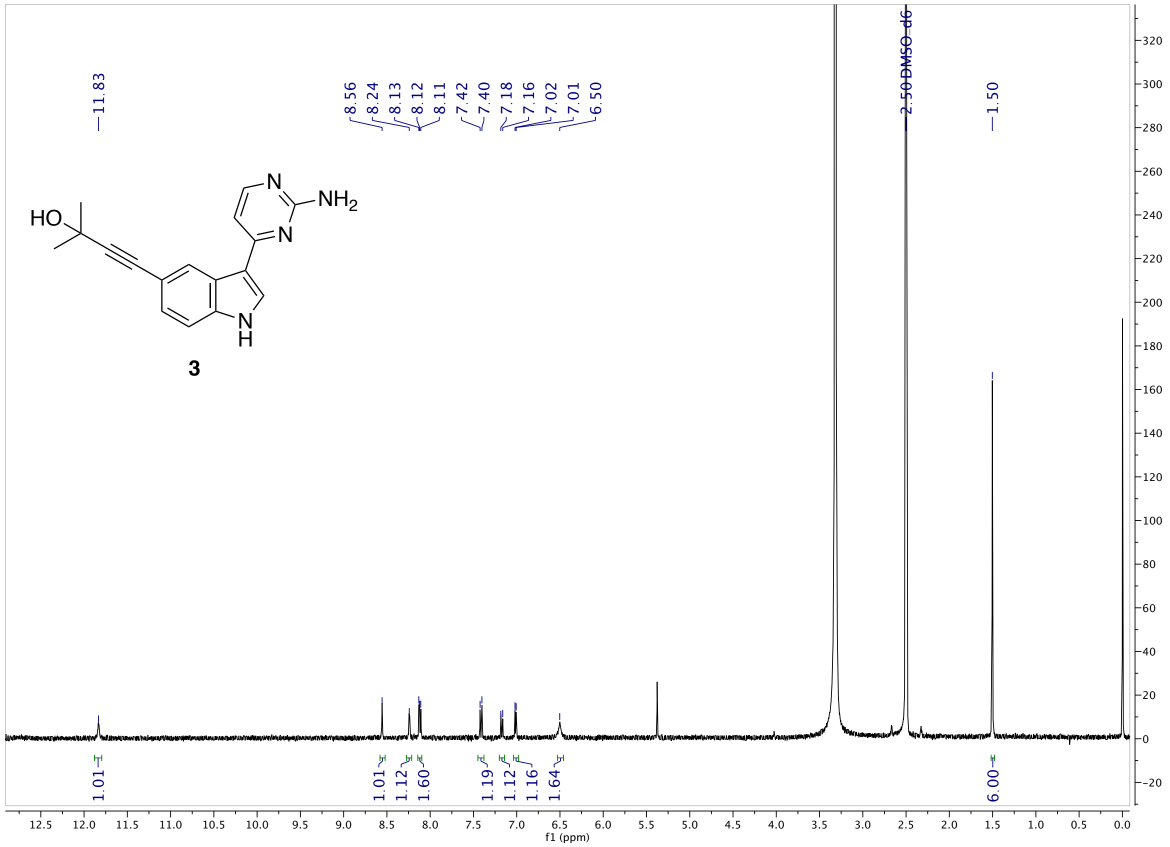

$^1$H NMR (400 MHz, DMSO-$d_6$) 1-(2-amino-5,6,7,8-tetrahydropyrimido[4',5':3,4]cyclohepta[1,2-b]indol-11-yl)-3-ethylpent-1-yn-3-ol (**9**):

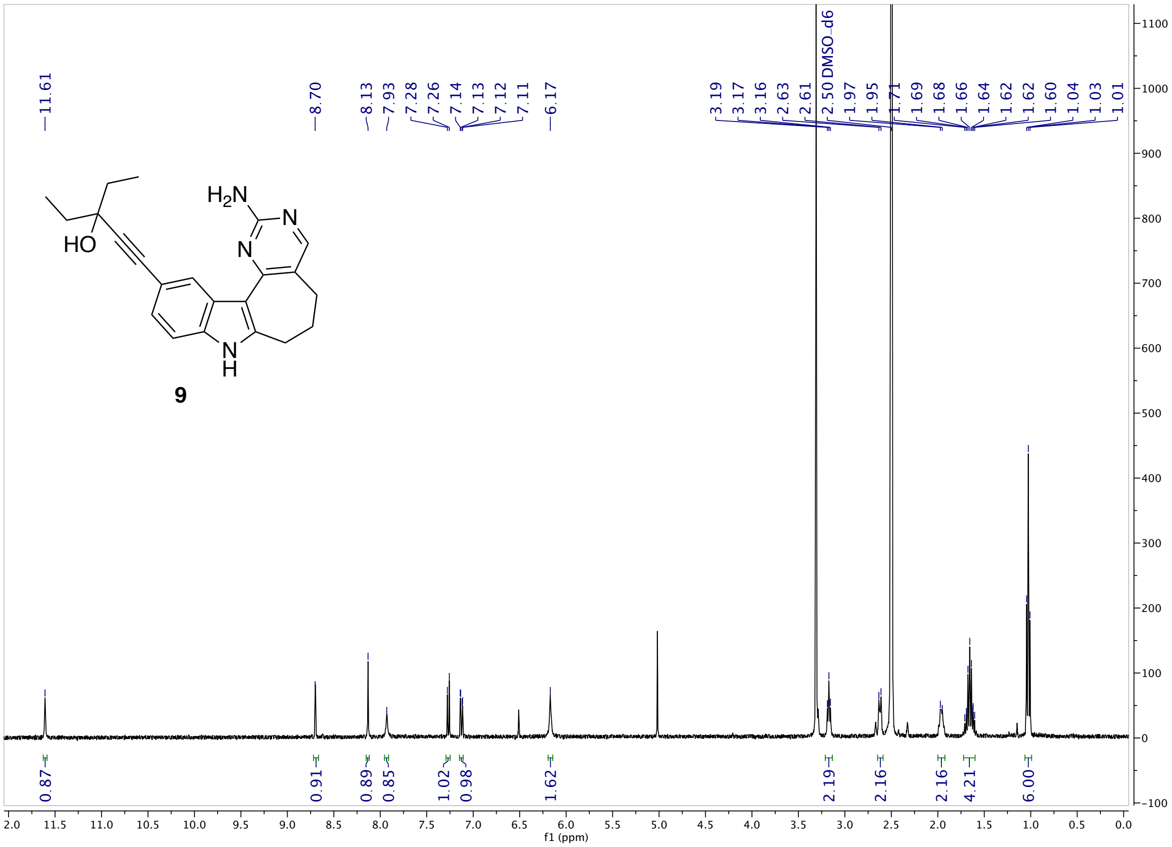

[1]H NMR (400 MHz, MeOD-*d₄*) 1-(2-amino-5,6,7,8-tetrahydropyrimido[4',5':3,4]cyclohepta[1,2-b]indol-11-yl)-3-methylpent-1-yn-3-ol (**10**):

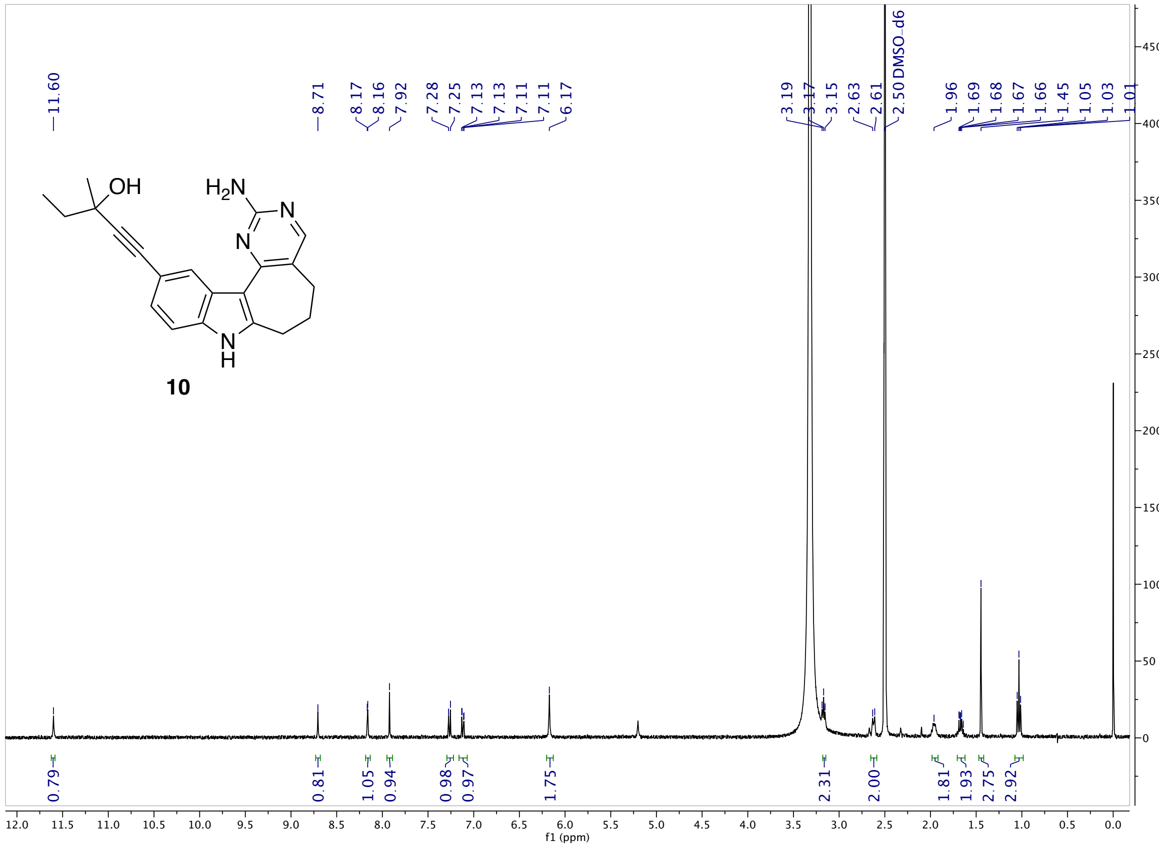

$^1$H NMR (400 MHz, MeOD-$d_4$) N-(3-(3-(2-aminopyrimidin-4-yl)-5-(3-hydroxy-3-methylbut-1-yn-1-yl)-1H-indol-1-yl)propyl)-3-(5,5-difluoro-7-(1H-pyrrol-2-yl)-5H-5λ4,6λ4-dipyrrolo[1,2-c:2',1'-f][1,3,2]diazaborinin-3-yl)propenamide (**29**):

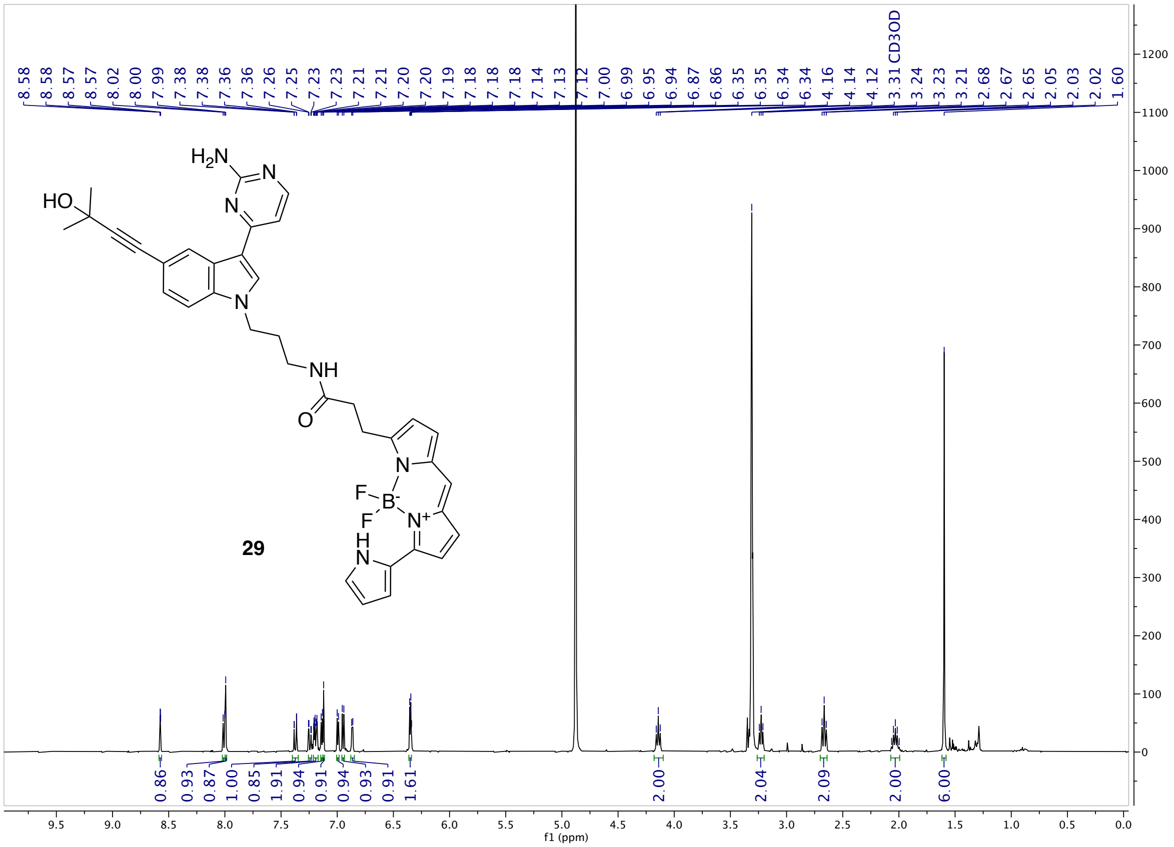
